# Phylogenetically distinct *Vibrio mediterranei* lineages confer robust protection under thermal stress against oyster pathogens

**DOI:** 10.64898/2026.02.09.704869

**Authors:** Steph Smith, Ami Wilbur, Jaypee S. Samson, Yesmarie De La Flor, Marta Gomez-Chiarri, Blake Ushijima, Rachel T. Noble

## Abstract

Marine bivalve mortality events cause substantial economic losses in aquaculture and threaten global food security. While pathogenic *Vibrio* species are frequently implicated, growing evidence suggests that loss of beneficial microbes can increase host susceptibility to disease. We previously observed that *Vibrio mediterranei* was consistently isolated from healthy oysters but systematically disappeared prior to mortality events, coinciding with proliferation of pathogenic *Vibrio* species. Here, we test whether this pattern reflects a protective functional role. Pre-colonization with *V. mediterranei* strain Vm02 increased *Crassostrea virginica* oyster larval survival from 10-19% (pathogen-only controls) to 94-97% when challenged with *V. harveyi* or *V. coralliilyticus*, representing near-complete protection from pathogen-induced mortality. Protection was maintained at both ambient (28°C) and thermal stress (32°C) temperatures where pathogen virulence is enhanced, rapid (effective from co-inoculation), and durable (maintained for >96 hours). Fluorescence microscopy confirmed stable colonization of larval digestive tissues by fluorescently-tagged Vm02. Larval colonization by eleven *V. mediterranei* strains revealed three distinct phenotypes – protective, pathogenic, and intermediate – corresponding to monophyletic clades with 97.1-97.8% average nucleotide identity between protective and pathogenic lineages. Pangenome analysis identified 230 protective-specific versus 80 pathogenic-specific orthogroups. Protective strains encode unique regulatory systems, stress tolerance mechanisms, and metabolic versatility while lacking Type I and Type VI secretion system variants associated with pathogenicity. Together, these findings demonstrate that beneficial versus pathogenic phenotypes are phylogenetically constrained within distinct *V. mediterranei* lineages. This supports reports that *V. mediterranei* acts as both a pathogen and potential symbiont in marine hosts and reveals a clade that provides robust protection against oyster pathogens.

**Importance:** Aquaculture disease management has traditionally emphasized either prophylactic treatment using antibiotics to avoid disease and dysbiosis or has focused entirely on pathogen detection. Both of these approaches have overlooked the potential contributions of beneficial microbes to host defense and grow-out performance. Developing beneficial probiotic tools for disease prevention represents an emerging opportunity for sustainable aquaculture management. This study demonstrates that specific lineages of *Vibrio mediterranei* function as protective symbionts capable of rescuing oyster larvae from near-complete pathogen-induced mortality. By integrating field observations of microbial succession during mortality events with experimental validation and comparative genomics, we show that protective versus pathogenic phenotypes are phylogenetically constrained within *V. mediterranei* clades separated by 97.1-97.8% average nucleotide identity. This resolution of strain-level functional variation provides fundamental insights into how host-microbe mutualisms evolve within species complexes that also harbor pathogens. The unique genomic markers identified here enable reliable screening for protective symbionts, while the temperature-stable and durable protection demonstrated in this study highlights the potential for biological control strategies in shellfish hatcheries increasingly affected by warming oceans and *Vibrio*-driven mortality events.

## 3 Introduction

Marine bivalve mortality events represent a major obstacle to the economic viability of aquaculture, causing annual losses in the billions of dollars annually and undermining the consistency and reliability of shellfish markets while threatening food security across coastal regions (Rato et al. 2022; Masanja et al. 2024; Smaal et al. 2019; *The State of Food Security and Nutrition in the World 2023* 2023). These events impact all major shellfish species and can result in stock losses exceeding 90%, with frequency and severity increasing due to ocean warming and environmental stress (Stewart-Sinclair et al. 2020; Ben-Horin et al. 2024; Matkarimov et al. 2025). Disease management efforts have largely focused on identifying and controlling pathogenic microorganisms, particularly *Vibrio* species, which are frequently implicated as causative agents of mortality across marine invertebrates (Richards et al. 2015; Bruto et al. 2017; Xu et al. 2022). Members of the Harveyi clade, including *V. harveyi* and *V. coralliilyticus*, are particularly problematic in shellfish hatcheries where they cause acute larval mortality through extracellular toxins and proteases (Zhang, He, and Austin 2020; Richards et al. 2015; Ushijima et al. 2022).

However, increasing evidence suggests that a pathogen-centric view overlooks critical features of host-microbe interactions that govern disease susceptibility. Across terrestrial, human, and marine systems, the depletion of beneficial microbes can predispose hosts to pathogenic colonization and contribute to dysbiosis-driven mortality (Zaneveld, McMinds, and Vega Thurber 2017; Egan and Gardiner 2016; Stevens, Bates, and King 2021). In Pacific oysters infected with OsHV-1, microbiome destabilization precedes mortality and involves depletion of diverse bacterial communities coupled with proliferation of opportunistic *Vibrio* species (Petton et al. 2021; De Lorgeril et al. 2018; Clerissi et al. 2020). Similar patterns occur in scallops exposed to harmful algal blooms, where paralytic shellfish toxins induce gut dysbiosis characterized by increased *Vibrio* abundance and reduced microbial diversity (Wei et al. 2024). These observations suggest that community composition, including the presence or absence of specific beneficial taxa, may ultimately determine disease trajectories in marine bivalves.

Our previous analysis of *Vibrio* communities in eastern oysters (*Crassostrea virginica*) revealed distinct assemblages associated with healthy versus diseased states (Smith et al. 2025). Most notably, *Vibrio mediterranei* was consistently isolated only in low-mortality periods and was systematically absent during mortality progression. The consistent loss of *V. mediterranei* associated with mortality raised the possibility that specific beneficial *Vibrios* may influence host resilience.

*Vibrio mediterranei* is ubiquitous across marine habitats, isolated from seawater, sediments, and diverse marine invertebrate hosts worldwide. Yet this species presents an ecological paradox: while some strains cause disease in corals (Kushmaro et al. 2001; Rubio-Portillo et al. 2020), pen shells (Andree et al. 2021), razor clams (Fan et al. 2023), and macroalgae (Yang et al. 2020), other strains are consistently recovered from healthy animals, including bacteriocin-producing isolates with probiotic potential (Serrano et al. 2018; Carraturo et al. 2006; Worden et al. 2022; Smith et al. 2025). Such dual ecological roles are well-established in other bacterial systems; *Staphylococcus aureus*, for example, exists as a harmless commensal in many humans while causing severe disease in others. These contradictory observations, together with the systematic disappearance of *V. mediterranei* prior to oyster mortality events (Smith et al. 2025), suggest that strain-level variation determines whether this species functions as a pathogen or beneficial symbiont.

Environmental temperature strongly influences *Vibrio* community composition and virulence (Kimes et al. 2012; Green et al. 2019; Vezzulli et al. 2016). Temperature-dependent virulence has been demonstrated in *V. mediterranei* infections of *Pinna nobilis* (Andree et al. 2021), and in coral systems where warming drives synergistic co-infection dynamics (Rubio-Portillo et al. 2020). Given that marine heatwaves are intensifying globally (Oliver et al. 2018), it is essential to understand how temperature affects both pathogenic and beneficial interactions between hosts and their associated microbiota.

Based on these observations, we hypothesized that: (1) *V. mediterranei* isolates consistently associated with healthy oysters function as protective symbionts rather than opportunistic colonizers; (2) protective versus pathogenic phenotypes represent phylogenetically constrained strategies within the *V. mediterranei* species complex; and (3) loss of protective *V. mediterranei* strains may create ecological conditions that facilitate pathogenic *Vibrio*-induced mortality.

Here, we test these hypotheses using controlled small-scale larval challenge experiments in which larval survival, bacterial colonization patterns, and physiological condition were monitored, combined with comparative genomics across strains isolated from healthy and diseased hosts. We show that specific *V. mediterranei* lineages provide rapid and robust protection against established oyster pathogens under thermal stress conditions, and that beneficial and pathogenic phenotypes are constrained within distinct evolutionary clades. These findings provide mechanistic insight into how mutualistic traits evolve within *Vibrio* species complexes and offer practical applications for disease forecasting and biological control in aquaculture systems.

## 4 Materials and Methods

### 4.1 Bacterial strains and culture conditions

All bacterial strains used in this study are listed in Table 1. Protective *Vibrio mediterranei* strains Vm02 (PNB23_20_7), PNB23_3_3, PNB22_9_1, PNB22_6_1, PNB22_7_2, and PNB22_2_1 were isolated in 2022-2023 from healthy adult eastern oysters (*Crassostrea virginica*) in North Carolina as described previously (Smith et al. 2025). Intermediate *V. mediterranei* strains PPP23_1_1 and PPP23_1_2 were isolated from healthy adult eastern oysters at a separate North Carolina commercial aquaculture farm during the same sampling period. Intermediate strains FYC25 and FYC26 were isolated by the University of Rhode Island from larvae collected from a mortality event at an eastern oyster hatchery in the Northeast US.

**Table 1.**
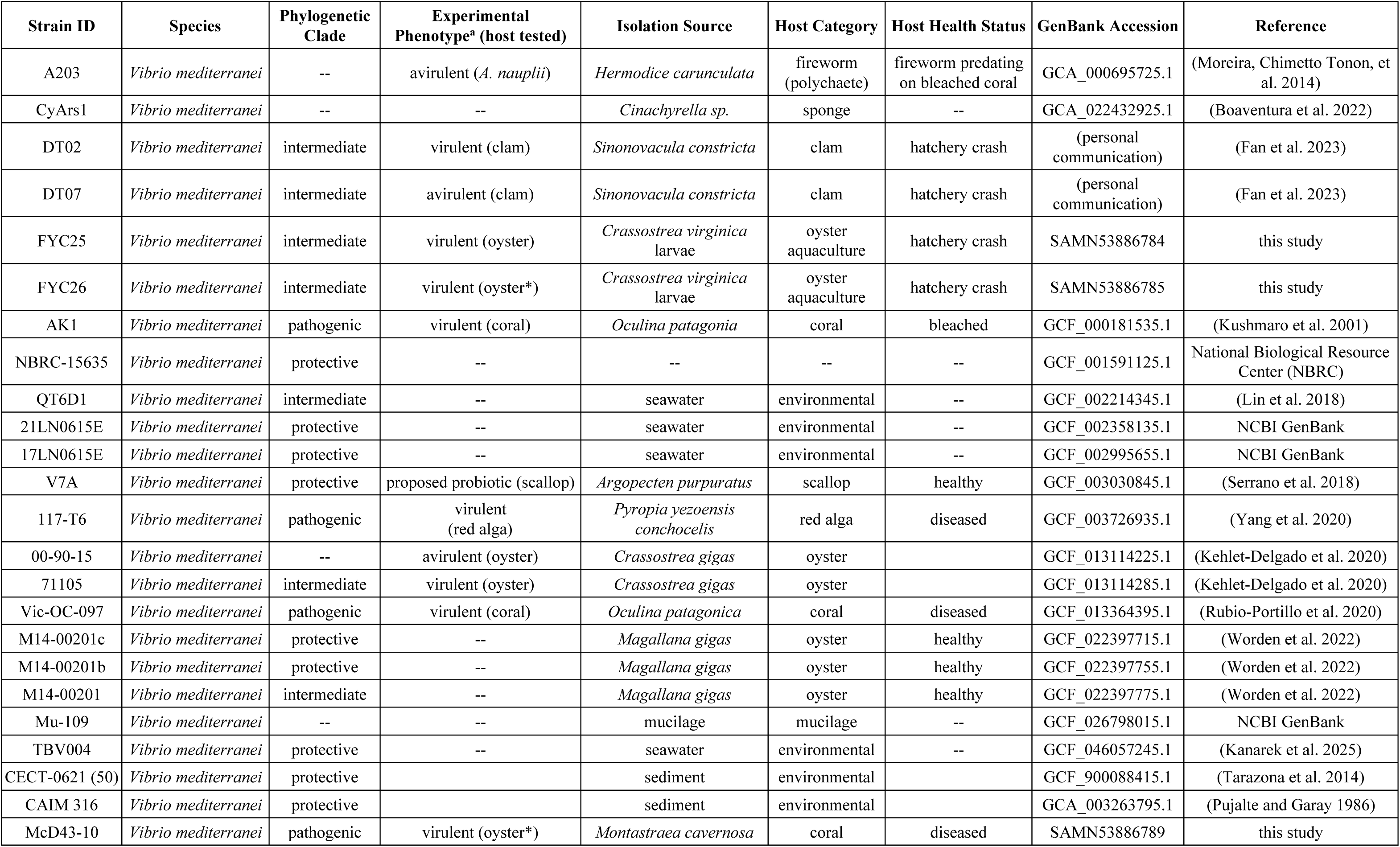
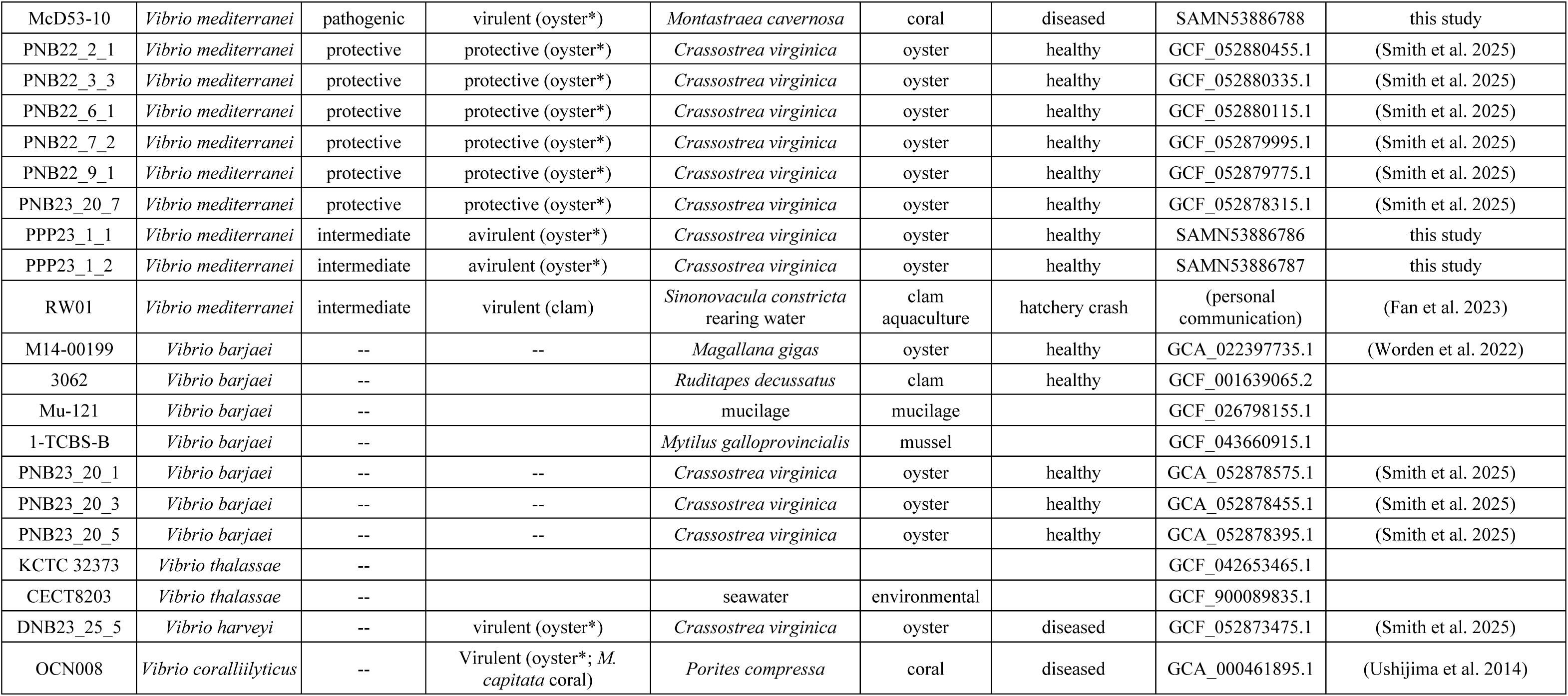
*Vibrio* strains used in this study. Complete table of all bacterial strains used in experimental assays and comparative genomic analyses. For experimentally tested strains (n=11), phenotypic classifications are based on larval challenge assays at 32°C (Figure 4). For reference *V. mediterranei* strains (n=22), phenotypic classifications are based on previous literature reports, including isolation host phenotype and/or subsequent experimental validation. Isolation sources and host health status provide ecological context for interpreting strain-phenotype relationships. ^a^Asterisk indicates phenotype was experimentally tested in this study.

| Strain ID | Species | Phylogenetic Clade | Experimental Phenotype<sup>a</sup> (host tested) | Isolation Source | Host Category | Host Health Status | GenBank Accession | Reference |
| --- | --- | --- | --- | --- | --- | --- | --- | --- |
| A203 | *Vibrio mediterranei* | -- | avirulent (*A. nauplii*) | *Hermodice carunculata* | fireworm (polychaete) | fireworm predating on bleached coral | GCA_000695725.1 | (Moreira, Chimetto Tonon, et al. 2014) |
| CyArs1 | *Vibrio mediterranei* | -- | -- | *Cinachyrella sp.* | sponge | -- | GCA_022432925.1 | (Boaventura et al. 2022) |
| DT02 | *Vibrio mediterranei* | intermediate | virulent (clam) | *Sinonovacula constricta* | clam | hatchery crash | (personal communication) | (Fan et al. 2023) |
| DT07 | *Vibrio mediterranei* | intermediate | avirulent (clam) | *Sinonovacula constricta* | clam | hatchery crash | (personal communication) | (Fan et al. 2023) |
| FYC25 | *Vibrio mediterranei* | intermediate | virulent (oyster) | *Crassostrea virginica* larvae | oyster aquaculture | hatchery crash | SAMN53886784 | this study |
| FYC26 | *Vibrio mediterranei* | intermediate | virulent (oyster*) | *Crassostrea virginica* larvae | oyster aquaculture | hatchery crash | SAMN53886785 | this study |
| AK1 | *Vibrio mediterranei* | pathogenic | virulent (coral) | *Oculina patagonica* | coral | bleached | GCF_000181535.1 | (Kushmaro et al. 2001) |
| NBRC-15635 | *Vibrio mediterranei* | protective | -- | -- | -- | -- | GCF_001591125.1 | National Biological Resource Center (NBRC) |
| QT6D1 | *Vibrio mediterranei* | intermediate | -- | seawater | environmental | -- | GCF_002214345.1 | (Lin et al. 2018) |
| 21LN0615E | *Vibrio mediterranei* | protective | -- | seawater | environmental | -- | GCF_002358135.1 | NCBI GenBank |
| 17LN0615E | *Vibrio mediterranei* | protective | -- | seawater | environmental | -- | GCF_002995655.1 | NCBI GenBank |
| V7A | *Vibrio mediterranei* | protective | proposed probiotic (scallop) | *Argopecten purpuratus* | scallop | healthy | GCF_003030845.1 | (Serrano et al. 2018) |
| 117-T6 | *Vibrio mediterranei* | pathogenic | virulent (red alga) | *Pyropia yezoensis conchocelis* | red alga | diseased | GCF_003726935.1 | (Yang et al. 2020) |
| 00-90-15 | *Vibrio mediterranei* | -- | avirulent (oyster) | *Crassostrea gigas* | oyster |  | GCF_013114225.1 | (Kehlet-Delgado et al. 2020) |
| 71105 | *Vibrio mediterranei* | intermediate | virulent (oyster) | *Crassostrea gigas* | oyster |  | GCF_013114285.1 | (Kehlet-Delgado et al. 2020) |
| Vic-OC-097 | *Vibrio mediterranei* | pathogenic | virulent (coral) | *Oculina patagonica* | coral | diseased | GCF_013364395.1 | (Rubio-Portillo et al. 2020) |
| M14-00201c | *Vibrio mediterranei* | protective | -- | *Magallana gigas* | oyster | healthy | GCF_022397715.1 | (Worden et al. 2022) |
| M14-00201b | *Vibrio mediterranei* | protective | -- | *Magallana gigas* | oyster | healthy | GCF_022397755.1 | (Worden et al. 2022) |
| M14-00201 | *Vibrio mediterranei* | intermediate | -- | *Magallana gigas* | oyster | healthy | GCF_022397775.1 | (Worden et al. 2022) |
| Mu-109 | *Vibrio mediterranei* | -- | -- | mucilage | mucilage | -- | GCF_026798015.1 | NCBI GenBank |
| TBV004 | *Vibrio mediterranei* | protective | -- | seawater | environmental | -- | GCF_046057245.1 | (Kanarek et al. 2025) |
| CECT-0621 (50) | *Vibrio mediterranei* | protective |  | sediment | environmental |  | GCF_900088415.1 | (Tarazona et al. 2014) |
| CAIM 316 | *Vibrio mediterranei* | protective |  | sediment | environmental |  | GCA_003263795.1 | (Pujalte and Garay 1986) |
| McD43-10 | *Vibrio mediterranei* | pathogenic | virulent (oyster*) | *Montastraea cavernosa* | coral | diseased | SAMN53886789 | this study |
| McD53-10 | *Vibrio mediterranei* | pathogenic | virulent (oyster*) | *Montastraea cavernosa* | coral | diseased | SAMN53886788 | this study |
| PNB22_2_1 | *Vibrio mediterranei* | protective | protective (oyster*) | *Crassostrea virginica* | oyster | healthy | GCF_052880455.1 | (Smith et al. 2025) |
| PNB22_3_3 | *Vibrio mediterranei* | protective | protective (oyster*) | *Crassostrea virginica* | oyster | healthy | GCF_052880335.1 | (Smith et al. 2025) |
| PNB22_6_1 | *Vibrio mediterranei* | protective | protective (oyster*) | *Crassostrea virginica* | oyster | healthy | GCF_052880115.1 | (Smith et al. 2025) |
| PNB22_7_2 | *Vibrio mediterranei* | protective | protective (oyster*) | *Crassostrea virginica* | oyster | healthy | GCF_052879995.1 | (Smith et al. 2025) |
| PNB22_9_1 | *Vibrio mediterranei* | protective | protective (oyster*) | *Crassostrea virginica* | oyster | healthy | GCF_052879775.1 | (Smith et al. 2025) |
| PNB23_20_7 | *Vibrio mediterranei* | protective | protective (oyster*) | *Crassostrea virginica* | oyster | healthy | GCF_052878315.1 | (Smith et al. 2025) |
| PPP23_1_1 | *Vibrio mediterranei* | intermediate | avirulent (oyster*) | *Crassostrea virginica* | oyster | healthy | SAMN53886786 | this study |
| PPP23_1_2 | *Vibrio mediterranei* | intermediate | avirulent (oyster*) | *Crassostrea virginica* | oyster | healthy | SAMN53886787 | this study |
| RW01 | *Vibrio mediterranei* | intermediate | virulent (clam) | *Sinonovacula constricta*<br>rearing water | clam<br>aquaculture | hatchery crash | (personal<br>communication) | (Fan et al. 2023) |
| M14-00199 | *Vibrio barjanei* | -- | -- | *Magallana gigas* | oyster | healthy | GCA_022397735.1 | (Worden et al. 2022) |
| 3062 | *Vibrio barjanei* | -- |  | *Ruditapes decussatus* | clam | healthy | GCF_001639065.2 |  |
| Mu-121 | *Vibrio barjanei* | -- |  | mucilage | mucilage |  | GCF_026798155.1 |  |
| 1-TCBS-B | *Vibrio barjanei* | -- |  | *Mytilus galloprovincialis* | mussel |  | GCF_043660915.1 |  |
| PNB23_20_1 | *Vibrio barjanei* | -- | -- | *Crassostrea virginica* | oyster | healthy | GCA_052878575.1 | (Smith et al. 2025) |
| PNB23_20_3 | *Vibrio barjanei* | -- | -- | *Crassostrea virginica* | oyster | healthy | GCA_052878455.1 | (Smith et al. 2025) |
| PNB23_20_5 | *Vibrio barjanei* | -- | -- | *Crassostrea virginica* | oyster | healthy | GCA_052878395.1 | (Smith et al. 2025) |
| KCTC 32373 | *Vibrio thalassae* | -- |  |  |  |  | GCF_042653465.1 |  |
| CECT8203 | *Vibrio thalassae* | -- |  | seawater | environmental |  | GCF_900089835.1 |  |
| DNB23_25_5 | *Vibrio harveyi* | -- | virulent (oyster*) | *Crassostrea virginica* | oyster | diseased | GCF_052873475.1 | (Smith et al. 2025) |
| OCN008 | *Vibrio coralliilyticus* | -- | Virulent (oyster*; *M. capitata* coral) | *Porites compressa* | coral | diseased | GCA_000461895.1 | (Ushijima et al. 2014) |

Pathogenic *V. mediterranei* strains McD53-9 and McD43-10 were isolated from *Montastraea cavernosa* corals displaying tissue loss lesions in the Florida Keys, USA (corals collected from Sand Key under permit issued by the Florida Keys National Marine Sanctuary [FKNMS-2017-128-A2]). Coral samples were collected and processed as previously described (Ushijima et al. 2020). However, initial coral tissue/mucus slurry samples were plated onto seawater agar (SWA) and incubated under anaerobic conditions at 28 ℃ within a BD GasPack EZ container system (Becton, Dickinson and Company) for five days. Colonies were streaked for purification on SWA before being incubated again anaerobically. Purified strains were grown in seawater broth (SWB) before being cryopreserved with 20% (v/v) glycerol at −80 ℃.

Pathogenic controls included *V. harveyi* Vh11 (DNB23_25_5), isolated from oysters during a 2023 mortality event (Smith et al. 2025), and *V. coralliilyticus* OCN008, previously characterized as a coral and oyster pathogen (Ushijima et al. 2014; 2022).

All strains were stored at −80°C in Marine Broth 2216 (MB2216; BD Difco) supplemented with 20% (v/v) glycerol. For experiments, frozen stocks were streaked onto MB2216 agar plates and incubated at 28°C for 24h. Single colonies were inoculated into 3 mL MB2216 broth and grown overnight (16-18 hours) at 28°C with shaking at 200 rpm. On the day of experimentation, overnight cultures were diluted 1:100 into fresh MB2216 and grown to mid-exponential phase (OD600 = 0.3-0.5, ∼3-4 hours) at experimental temperature (28°C or 32°C). Cells were harvested by centrifugation (7,000 × g, 3 min, room temperature), washed twice in sterile artificial seawater (ASW; Instant Ocean, 30 ppt), and resuspended to 1×108 CFU mL-1 based on optical density.

### 4.2 Construction of GFP-expressing *V. mediterranei* Vm02

For colonization imaging, *V. mediterranei* Vm02 was transformed with plasmid pVSV102 (Dunn et al. 2006), which constitutively expresses green fluorescent protein (GFP) and confers kanamycin resistance. Transformation was performed by tri-parental conjugative mating using *Escherichia coli* S17-1 λpir as the donor. Donor and recipient cells were grown to mid-exponential phase, mixed at a 1:1 ratio with helper plasmid pEVS104 (Dunn et al. 2006), pelleted by centrifugation, and spotted onto MB2216 agar plates. After 8 hours incubation at 28°C, cells were resuspended in MB2216 broth and plated on LBS agar supplemented with 50 µg mL-1 kanamycin. Fluorescent colonies were confirmed by epifluorescence microscopy and maintained on MB2216 agar containing 50 µg mL-1 kanamycin.

### 4.3 Larval oyster culture

Eastern oyster (*C. virginica*) larvae were obtained from the Shellfish Research Hatchery (SRH; University of North Carolina Wilmington, NC, USA) and maintained following standard hatchery procedures. Larvae were reared in filtered, UV-treated water obtained from the hatchery at 28°C and ∼30 ppt with gentle aeration. Algal diets consisted of a mixed diet (*Chaetoceros mulleri, C. neogracilis, Isochrysis lutea,* and *Pavlova lutheri*) supplied at rations following Helm et al. 2004 (Helm et al. 2004).

For challenge experiments, veliger stage larvae (5-7 days post-fertilization) were concentrated by gentle sieving and resuspended in rearing water to a working density of ∼20 larvae mL-1. Larvae were distributed into sterile 50mL polystyrene tubes at ∼400 larvae per replicate volume (20 mL) and acclimated at the target experimental temperature (28°C or 32°C) for 1h prior to bacterial inoculation.

### 4.4 Larval challenge experiments

#### 4.4.1 Single-strain exposure and temperature dependence

To compare baseline virulence of protective, intermediate, and pathogenic *V. mediterranei* strains and control pathogens, larvae were exposed to individual strains at a defined cell density. Mid-exponential phase cultures were washed and resuspended in ASW as described above, then added to larval suspensions to achieve final concentrations of 106 CFU mL-1. Experiments were conducted at 28°C (ambient hatchery temperature) and 32°C (elevated temperature representing thermal stress).

Treatments included: (1) non-inoculated controls (larvae and algal feed only), (2) exposure to individual *V. mediterranei* strains, and (3) exposure to pathogenic controls Vh11 and OCN008. Larvae were incubated for 24h post-exposure before survival was assessed.

#### 4.4.2 Protection assays and pre-colonization experimental design

To test whether Vm02 confers protection against pathogenic *Vibrio*, larvae were exposed to combinations of Vm02 with Vh11 or OCN008. Each strain was added to a final concentration of 106 CFU mL-1. Experimental treatments included: (1) non-inoculated control (algae only), (2) Vm02 only, (3) pathogen only, and (4) combined Vm02 + pathogen.

Pre-colonization experiments tested the dependence of protection on colonization timing. Larvae were first inoculated with Vm02 at 106 CFU mL-1, incubated at 28°C, and then challenged with Vh11 at 106 CFU mL-1 after 0, 24, 48, or 96 h. Pre-colonization treatments were maintained at 28°C with a single Vm02 and Vh11 inoculation per treatment. After pathogen addition, larvae were incubated an additional 24 hours before survival assessment.

Co-inoculation evaluated the rapidity of protection: Vm02 and pathogen were added simultaneously at 106 CFU mL-1 each, and survival was measured after 24h.

All larval challenge experiments included at least two independent biological replicates (separate tubes) per treatment, and each experiment was repeated with larvae from two independent spawning cohorts. Experiments were conducted in temperature-controlled incubators set to the target temperature.

### 4.5 Larval survival assessment and relative larval survival (RLS)

Larval survival was quantified at 24h after the final bacterial exposure using light microscopy. The full volume of each replicate tube was gently mixed and transferred into a sterile Petri dish. Larvae were scored as live or dead based on established criteria: live larvae exhibited active velar ciliary beating, coordinated swimming, and intact shell and tissue morphology; dead larvae displayed complete absence of movement, tissue disintegration, or empty shells. All counts were performed by the same observer to ensure consistency.

Relative larval survival (RLS) was calculated to normalize for background mortality in controls:

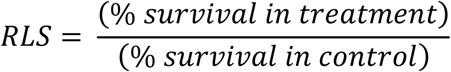

Where % survival = (number live larvae / total larvae counted) x 100. For protection experiments, RLS values were calculated relative to non-inoculated (larvae and algae only) controls. RLS >1.0 indicates survival enhanced above control levels; RLS = 1 indicates no effect; RLS<1.0 indicates mortality exceeding background.

### 4.6 Statistical analyses

All statistical analyses were performed in R version 4.3.1 using the tidyverse, rstatix, and ggpubr packages. Relative larval survival (RLS) was calculated for each replicate tube and served as the response variable for all statistical comparisons.

For baseline virulence comparisons (Figure 1), temperature effects on RLS were assessed independently for each bacterial strain using two-sample t-tests comparing 28°C versus 32°C treatments.

**Figure 1.**
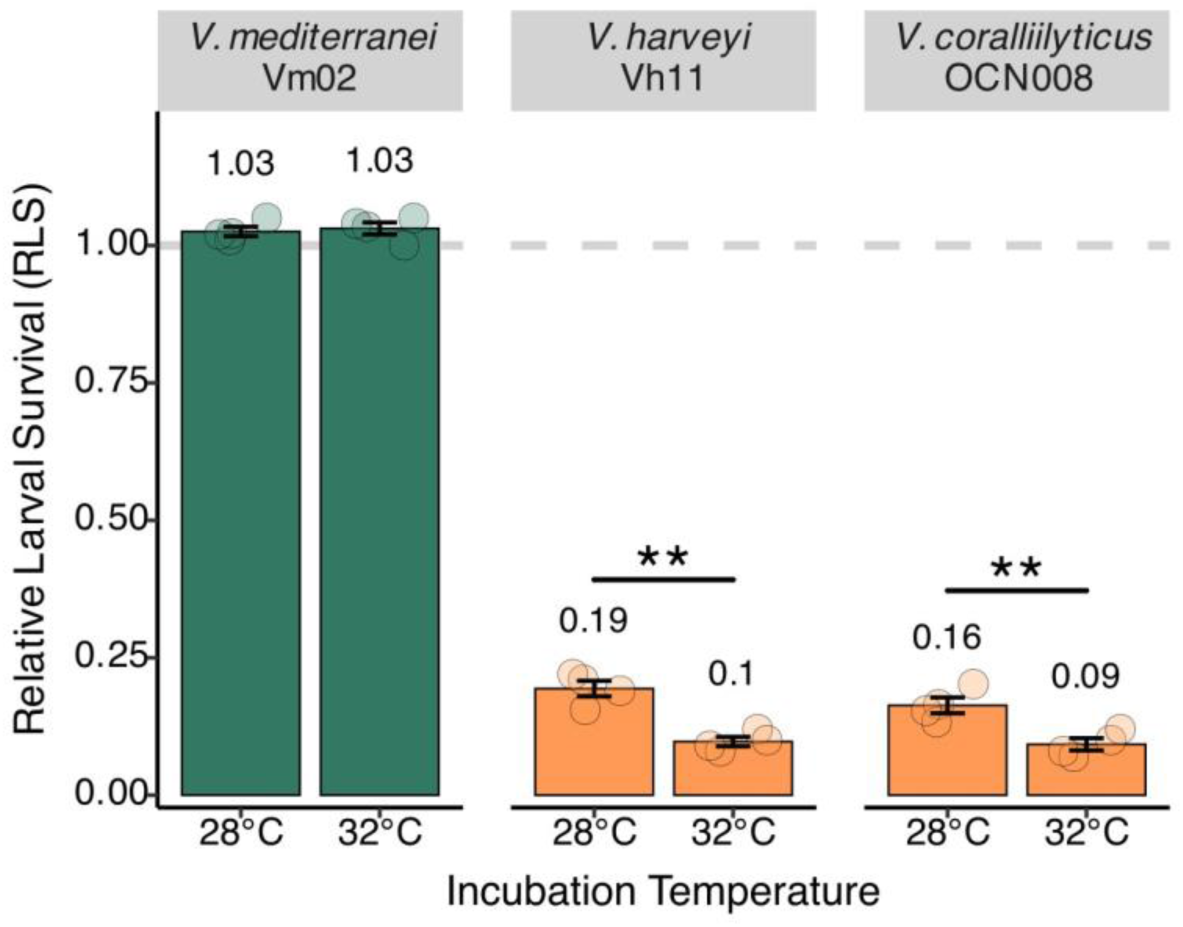
Strain-specific virulence of *Vibrio* species in oyster larvae. Eastern oyster larvae (*C. virginica*) exposed to *V. mediterranei* Vm02 show no mortality while pathogenic species *V. harveyi* Vh11 and *V. coralliilyticus* OCN008 cause severe temperature-enhanced mortality. Relative larval survival (RLS) following a 24-hour bacterial exposure at 28°C (ambient) or 32°C (thermal stress). Values shown as mean ± SE from four replicate tubes per treatment (2 spawning cohorts x 2 tubes per cohort; ∼400 larvae per tube). Data points represent individual tube measurements. **p<0.01 (two-sample t-tests comparing temperatures within each strain.

For protection experiments testing multiple conditions (Figure 2B), differences between pathogen-only and Vm02-protected treatments were evaluated using two-sample t-tests performed independently within each pathogen x temperature combination. P-values were adjusted for multiple comparisons across the four pathogen-temperature conditions using the Benjamini-Hochberg false discovery rate (FDR) method.

**Figure 2.**
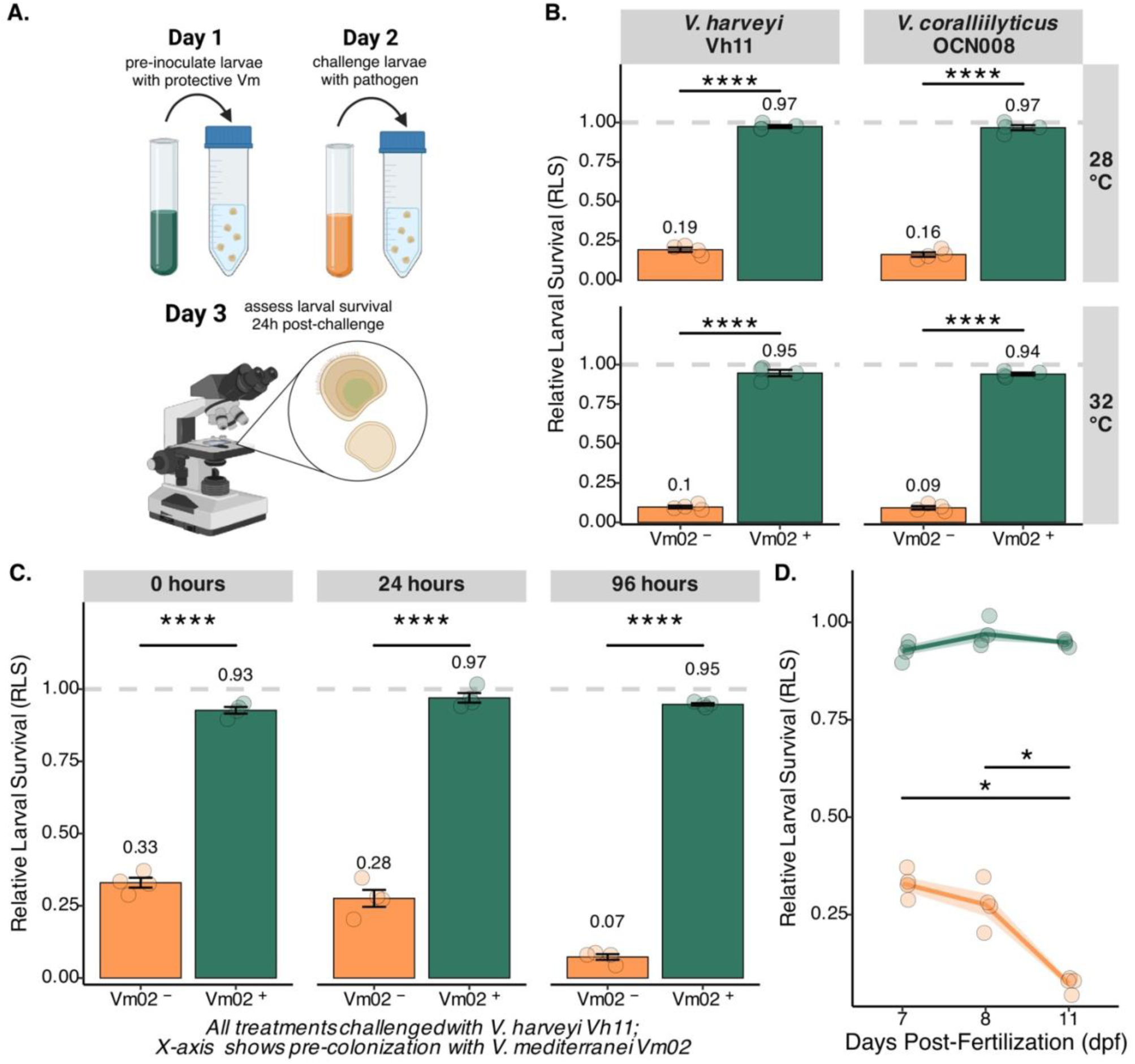
*V. mediterranei* Vm02 pre-colonization provides robust protection against pathogenic *Vibrio* species. **(A)** Experimental timeline showing pre-colonization and challenge protocol. **(B)** Robust protection against *V. harveyi* Vh11 and *V. coralliilyticus* OCN008 at 28°C and 32°C. Pre-colonization rescues larvae from ∼10-19% to 94-97% survival. **(C)** Protection efficacy across different pre-colonization intervals (0-96 hours). **(D)** Age-dependent susceptibility and protection across larval development (7-11 dpf). Values shown as mean ± SE from four replicate tubes per treatment (2 spawning cohorts x2 tubes per cohort; ∼400 larvae per tube). Data points represent individual tube measurements. Statistical comparisons: **(B)** two-sample t-tests with Benjamini-Hochberg FDR correction; **(C)** two-sample t-tests, uncorrected, **(D)** Kruskal-Wallis test with pairwise Wilcoxon tests and FDR correction. ****p<0.0001, *p<0.05.

For pre-colonization timing experiment (Figure 2C), protection efficacy was compared between Vm02-protected and unprotected larvae using two-sample t-tests performed independently at each pre-colonization interval (0h, 24h, 96h).

For age-dependent susceptibility experiments (Figure 2D), differences in larval survival across developmental stages were assessed separately for protection and evaluated whether survival differed across larval ages (7, 8, and 11 days post-fertilization). When significant (p<0.05), pairwise comparisons between age groups were conducted using Wilcoxon rank-sum tests with Benjamini-Hochberg FDR correction.

All experiments included two independent biological replicates (separate spawning cohorts) with two technical replicates (tubes) per treatment per cohort, for a total of four replicate tubes per treatment. Approximately 400 larvae were assessed per tube. Data are presented as mean ± standard error (SE) calculated across replicate tubes. Statistical significance was defined as α = 0.05 for all tests.

### 4.7 Fluorescence microscopy of larval colonization

To visualize bacterial colonization, larvae were inoculated with GFP-expressing Vm02 using the same inoculation schemes described for challenge experiments. At 24h and 96h post-inoculation, ∼1mL of larval suspension (∼20 larvae) per replicate was fixed in 4% paraformaldehyde (PFA) in PBS and stored at 4°C until imaging.

Immediately before imaging, fixed larvae were transferred onto glass microscope slides and immobilized under 2% (w/v) agarose pads prepared in sterile ASW adjusted to 20 psu. Coverslips were applied and slides were imaged immediately.

Fluorescence microscopy was performed using a Nikon Eclipse E600 upright microscope equipped with a Nikon DS-Fi3 camera and Sola Lumencor light engine. Images were acquired using a Nikon CFI Plan Fluor 40X objective with standardized GFP filter sets. Transmitted light and fluorescence images were captured sequentially for each field of view to visualize both overall morphology and bacterial colonization patterns. Exposure time, gain, and illumination settings were kept constant within each experiment. Image processing was performed using FIJI/ImageJ v1.53 (Schindelin et al. 2012). Brightness and contrast adjustments were applied uniformly across all images; no other manipulations were performed.

### 4.8 Genome sequencing and assembly

Whole-genome sequencing was performed on six *V. mediterranei* strains not previously characterized: pathogenic strains McD43-10 and McD53-9, and intermediate strains FYC25, FYC26, PPP23_1_1, and PPP23_1_2. All other *V. mediterranei* genomes used in comparative analyses were obtained from our previous study (Smith et al. 2025) or obtained from NCBI GenBank (accession numbers in Table 1).

Long-read sequencing was performed at SeqCoast Genomics on a PromethION 2 platform (Oxford Nanopore Technologies) (FYC26, PPP23_1_1, PPP23_1_2, McD53-9, McD43-10), or at the University of Rhode Island on a MinION Mk1B (Oxford Nanopore Technologies) (FYC25). Raw reads were trimmed using PoreChop and assembled using Unicycler with the nf-core/bacass v2.5.0 workflow (Daniel VM et al. 2025). Assembly quality metrics were assessed with QUAST. Draft genomes were deposited under BioProject accession PRJNA1379678, with individual assembly accession identifiers listed in Table 1.

### 4.9 Genome annotation, ANI, and phylogenomic analysis

Draft genomes, including reference assemblies retrieved from NCBI GenBank, were annotated using Prokka v1.14.6 (Seemann 2014) with consistent default parameters to ensure uniform gene prediction across the dataset.

Pangenome analysis and core genome alignment were performed using Panaroo v1.3.4 (Tonkin-Hill et al. 2020) with ‘—clean-mode strict’ to minimize spurious gene calls from assembly errors or potential contamination. The core genome was defined as genes present in ≥99% of all input genomes, yielding 3,684 core genes. Panaroo generated a multi-FASTA alignment of concatenated core genes that was used for phylogenomic reconstruction.

Maximum likelihood phylogenetic trees were constructed using IQ-TREE v2.2.0 (Minh et al. 2020). The best-fit substitution model was automatically selected using ModelFinder Plus (Kalyaanamoorthy et al. 2017) with Bayesian Information Criterion, identifying GTR+F+I+G4 as the optimal model. Phylogenetic inference was performed with 1,000 bootstrap replicates and 1,000 SH-aLRT tests to assess branch support. Trees were visualized with TreeViewer (Bianchini and Sánchez-Baracaldo 2024).

Pairwise average nucleotide identity (ANI) was calculated using pyANI v0.2.12 (Pritchard et al. 2016) with the ANIm method, which uses MUMmer (Marçais et al. 2018) for whole-genome alignment. Default parameters were used (minimum alignment length 1,000 bp, minimum alignment identity 30%, minimum alignment coverage 50%). Pairwise ANIm values were organized into a similarity matrix and visualized as a heatmap using pheatmap v1.0.12 in R with hierarchical clustering based on complete linkage. Within-clade and between-clade ANI distributions were summarized to quantify genomic divergence corresponding to phenotypic classifications.

### 4.10 Pangenome and functional analyses

Orthogroup classifications from Panaroo were used to characterize the pangenome structure of *V. mediterranei*. Clade-specific orthogroups were identified using custom Python scripts (Python v3.9.7) that parsed the Panaroo gene presence-absence matrix. Analysis was restricted to *V. mediterranei* strains specifically ascribed a host phenotype, either experimentally in this study or in the literature (colored tips Figure 5; Table 1). Protective-specific orthogroups were defined as present in 90% of experimentally-validated protective strains (n=6) and absent from 90% of pathogenic (n=7), intermediate-virulent (n=4), and intermediate-avirulent (n=6) strains. Pathogenic-specific, intermediate-virulent-specific, and intermediate-avirulent-specific orthogroups were defined analogously. This criterion (90% presence within-clade, 10% presence between-clades) ensured identification of genes consistently associated with each phenotypic group.

Functional annotation of all clade-specific protein sequences was performed using eggNOG-mapper v2.19 (Cantalapiedra et al. 2021) against the eggNOG v5.0 database with default parameters. This analysis assigned Clusters of Orthologous Groups (COG) functional categories, KEGG Orthology (KO identifiers), Gene Ontology (GO) terms, and PFAM domains to each protein. Orthogroups were assigned functional annotations based on their representative sequences.

Phylogenetic clade designations were assigned based on experimental phenotypic validation. The protective clade includes strains that demonstrated avirulence or enhanced survival (RLS≥1.0) when tested experimentally against bivalve larvae in this study or previous reports. The pathogenic clade includes only strains experimentally confirmed as pathogens in marine invertebrates. Strains from the same broader phylogenetic group but isolated from healthy hosts (A203 from healthy fireworm, CyArs1 from healthy sponge, Mu-109 from mucilage) are excluded from the pathogenic clade designation. Strain 00-90-15, which enhanced oyster larval survival experimentally (Kehlet-Delgado, Häse, and Mueller 2020) but falls phylogenetically between clades, was not assigned to any specific group.

## 5 Results

### 5.1 *V. mediterranei* Vm02 from healthy oysters is avirulent to larvae

To establish baseline virulence phenotypes of *Vibrio* strains isolated during our longitudinal mortality monitoring, we challenged 5-7 day post-spawn eastern oyster (*Crassostrea virginica*) larvae with representative *Vibrio* isolates at 106 CFU mL-1 under standardized conditions. *V. mediterranei* strain Vm02 (PNB23_20_7), isolated from a healthy adult oyster in the weeks preceding a mortality event, demonstrated complete avirulence to larval oysters across temperature conditions (Figure 1). At ambient hatchery temperature (28°C), Vm02-exposed larvae maintained relative larval survival (RLS) of 1.03 ± 0.02 compared to non-inoculated controls, indicating not only absence of pathogenicity but a modest enhancement of survival. This avirulent phenotype remained stable at elevated temperature (32°C), with RLS of 1.03 ± 0.01, demonstrating that elevated temperature does not induce virulence in this protective strain.

### 5.2 Temperature-dependent pathogen virulence

*V. harveyi* Vh11, isolated from an oyster several weeks later during a mass mortality event, caused severe mortality at both temperatures tested (RLS = 0.19 ± 0.04 at 28°C; 0.10 ± 0.02 at 32°C; p<0.01), representing 81% mortality above background levels. Elevated temperature (32°C) exacerbated pathogenicity, with RLS declining to 0.10 ± 0.02 (90% excess mortality). Similarly, *V. coralliilyticus* strain OCN008, a known coral pathogen in the Pacific and previously shown to cause mortality in Pacific oyster larvae (Ushijima et al. 2018), proved highly pathogenic to eastern oyster larvae. OCN008 caused comparable mortality to Vh11, with RLS of 0.16 ± 0.03 at 28°C and 0.09 ± 0.02 at 32°C. The temperature-independent avirulence of Vm02 contrasted sharply with the temperature-enhanced virulence of both pathogenic species, indicating a markedly different phenotype from pathogenic *Vibrios*.

### 5.3 *V. mediterranei* Vm02 pre-colonization protects against pathogens

Given the avirulent phenotype of Vm02, we investigated whether pre-colonization could protect larvae against subsequent pathogen challenge. Using a sequential inoculation design, larvae were pre-colonized with a single dose of Vm02 24 hours before challenge with either *V. harveyi* Vh11 or *V. coralliilyticus* OCN008 (Figure 2A). *V. mediterranei* pre-colonization dramatically altered infection outcomes at both temperature conditions (Figure 2B). At 28°C, larvae pre-colonized with Vm02 and subsequently challenged with Vh11 maintained 0.97 ± 0.02 RLS, compared to only 0.19 ± 0.04 survival in larvae exposed to Vh11 alone (p<0.0001, two-sample t-test, FDR-adjusted). This represents an 82% rescue effect, effectively restoring larval survival to control levels. Protection remained robust at 32°C, with pre-colonized larvae achieving 0.95 ± 0.02 RLS versus 0.10 ± 0.02 in unprotected controls (p<0.0001, two-sample t-test, FDR-adjusted), demonstrating that protective efficacy is maintained despite thermal stress conditions that enhance pathogen virulence.

Protection against *V. coralliilyticus* OCN008 proved equally effective. Pre-colonization with Vm02 increased survival from 0.16 ± 0.03 to 0.97 ± 0.02 at 28°C (p<0.0001, two-sample t-test, FDR-adjusted) and from 0.09 ± 0.02 to 0.94 ± 0.03 at 32°C (p<0.0001, two-sample t-test, FDR-adjusted). The consistency of protection across two phylogenetically distinct pathogens demonstrates that Vm02-mediated protection is not limited to a single pathogen species.

To test the effect of colonization timing and stability on prediction, we varied the time interval between Vm02 pre-colonization dose and Vh11 pathogen challenge. Each interval represents an independent treatment, with a single dose of Vm02 and a single subsequent dose of Vh11. Protection proved remarkably durable across all time intervals tested (Figure 2C). Even with co-inoculation (0h) when larvae were exposed to protector and pathogen simultaneously, Vm02 provided substantial protection, increasing RLS from 0.33 ± 0.05 in Vh11-only controls to 0.93 ± 0.03 in co-inoculated larvae (p<0.0001, two-sample t-test). Standard 24-hour pre-colonization yielded near-complete protection (RLS = 0.97 ± 0.02, p<0.0001), while extended pre-colonization periods of 96 hours maintained similarly high protection levels (RLS = 0.95 ± 0.03, p<0.0001). The rapid establishment and stability of protection suggested persistent colonization rather than transient effects. All time points assessed (up to 96 hours post-inoculation, corresponding to 11 dpf) occurred prior to larval metamorphosis.

Vm02 protection remained effective across early larval developmental stages, even as unprotected larvae became progressively more susceptible to Vh11 pathogen challenge with age (Figure 2D). We tested larvae at different ages pre-competence following a standard pre-colonization with Vm02. At 7 days post-fertilization (dpf), unprotected larvae maintained 0.35 ± 0.05 RLS when challenged with Vh11. This susceptibility increased significantly by 8 dpf (0.28 ± 0.08 RLS) and reached maximum vulnerability at 11 dpf (0.07 ± 0.02 RLS; Kruskal-Wallis χ²=6.8, p=0.033; pairwise Wilcoxon test with FDR correction: 7 vs 11 dpf=0.038). In contrast, Vm02-protected larvae maintained consistently high survival (>0.90 RLS) regardless of developmental stage (Kruskal-Wallis χ²=0.5, p=0.78), showing that protection remained robust across early larval stages.

### 5.4 *V. mediterranei* Vm02 establishes stable colonization of larval tissues

Fluorescence microscopy confirmed that protective Vm02 rapidly colonizes larval digestive tissues within 24 hours and maintains stable associations through at least 96 hours (Figure 3). To visualize colonization dynamics, we transformed Vm02 with a stable GFP-expressing plasmid (pVSV102). GFP-expressing Vm02 cells were consistently observed in association with the larval digestive system, with particularly dense colonization of the stomach and digestive diverticula. Colonization also extended to the velum, the primary feeding and swimming organ of larvae. In contrast, larvae exposed to *V. harveyi* Vh11 showed rapid tissue degradation, damaged cilia, and mortality within 24 hours.

**Figure 3.**
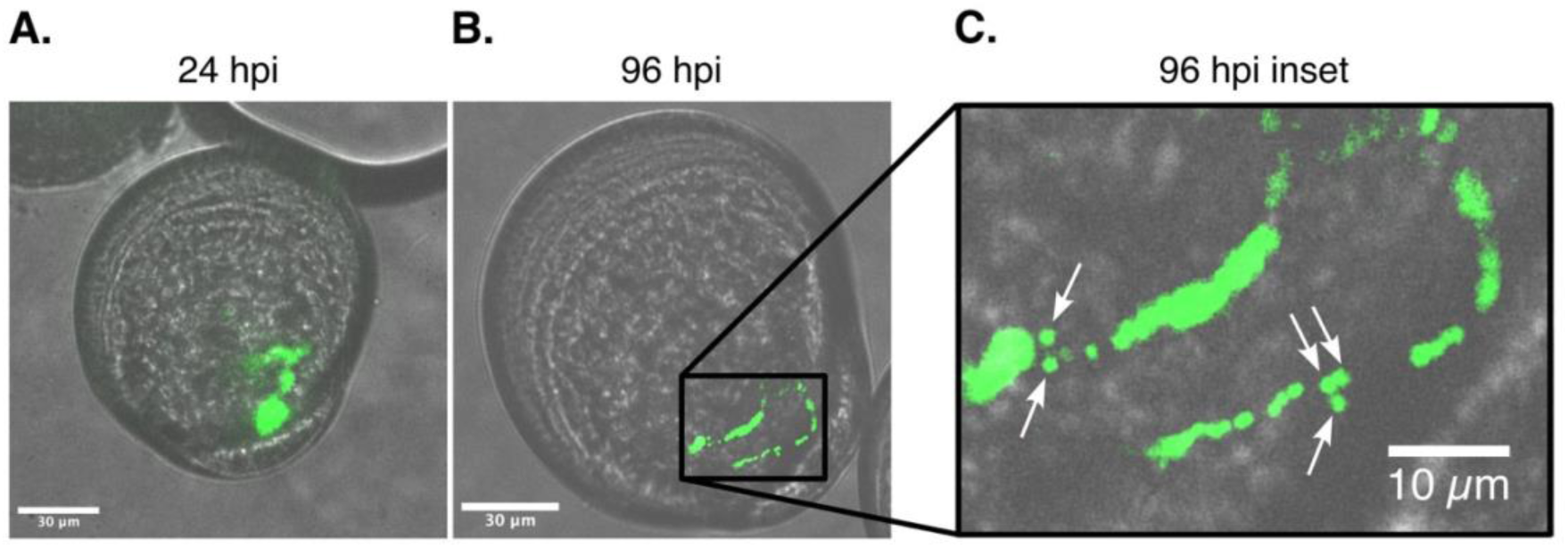
Stable colonization of oyster larvae by GFP-tagged *V. mediterranei* Vm02. Fluorescence microscopy showing persistent bacterial colonization at **(A)** 24 hours and **(B)** 96 hours post-inoculation. Green fluorescence indicates GFP-expressing Vm02 concentrated in digestive tissues and velum. **(C)** 96-hour inset showing individual Vm02 bacterial cells (arrows). Scale bars: 30µm **(A,B)**, 10µm **(C)**. Images representative of >20 larvae per timepoint repeated across two independent experiments.

The colonization pattern established by 24 hours persisted through at least 96 hours post-inoculation, with GFP-expressing Vm02 cells maintaining their association with digestive tissues. This persistent colonization corresponds with the sustained protection observed in our challenge experiments, indicating a durable physical association with larval digestive tissues.

### 5.5 *V. mediterranei* strains exhibit source-dependent virulence

To determine whether the protective phenotype was unique to strain Vm02 or characteristic of *V. mediterranei* more broadly, we tested eleven strains spanning the known diversity of this species complex (Table 1). All strains were tested clonally at 32°C to account for potential temperature-mediated virulence previously reported in *V. mediterranei* interactions with other marine hosts. Larval challenge assays revealed three distinct phenotypic categories that corresponded closely with strain isolation sources (Figure 4).

**Figure 4.**
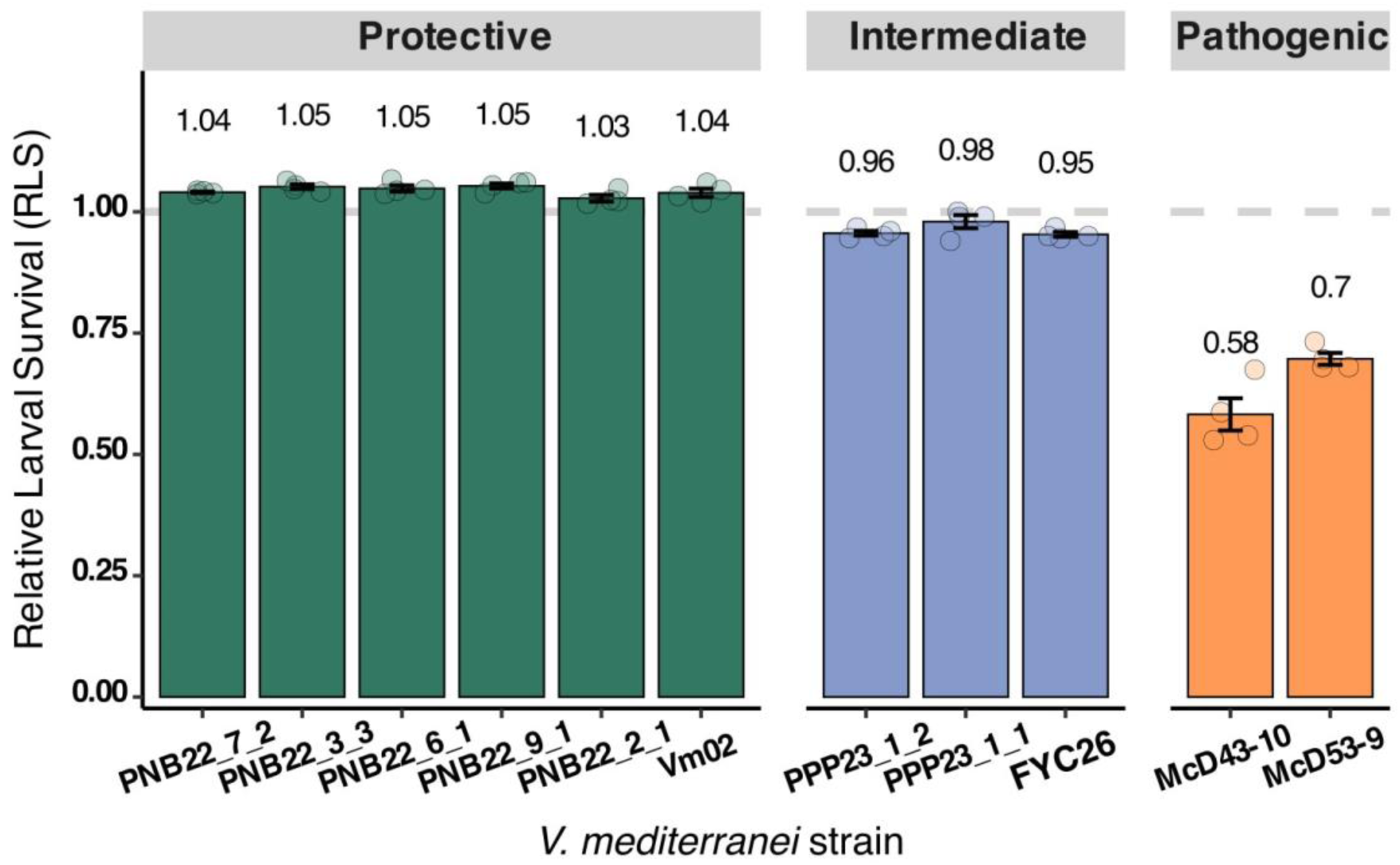
Strain-specific virulence phenotypes correspond to *V. mediterranei* isolation source. Eleven *V. mediterranei* strains exhibit three distinct phenotypes when tested against oyster larvae at 32°C. Protective strains (teal, n=6) from healthy oysters show RLS ≥1.0; intermediate strains (blue, n=3) show minimal virulence (RLS 0.95-0.98); pathogenic strains (orange, n=2) from diseased coral cause significant mortality (RLS 0.58-0.70). Values shown as mean ± SE from four replicate tubes per strain (2 spawning cohorts x 2 tubes per cohort; ∼400 larvae per tube). Data points represent individual tube measurements.

Six strains isolated from healthy adult oysters during low mortality periods demonstrated protective phenotypes (RLS=1.03-1.05), including our reference strain Vm02 (PNB23_20_7) and five additional isolates (PNB22_7_2, PNB22_2_2, PNB22_3_3, PNB22_6_1, PNB22_9_1). The consistency of the protective phenotype across multiple independent isolates from healthy oysters, collected across different sampling periods and locations, confirmed that protective capacity is a reproducible characteristic of oyster-associated *V. mediterranei* populations.

Two strains isolated from diseased coral tissues, McD53-9 and McD43-10, proved highly pathogenic to oyster larvae, causing 42% and 30% mortality respectively (RLS = 0.58 ± 0.03 and 0.70 ± 0.05). These strains were originally isolated from *Montastraea cavernosa* showing signs of tissue loss disease in the Florida Keys.

Three strains displayed intermediate virulence with slight but statistically significant larval mortality compared to non-inoculated controls. These included PPP23_1_2 (RLS = 0.96 ± 0.02), PPP23_1_1 (RLS = 0.98 ± 0.01), and Fyc26 (RLS = 0.95 ± 0.03). PPP23_1_2 and PPP23_1_2 were isolated from healthy adult oysters during the same sampling period as Vm02, although at a different farm, and FYC26 was isolated from larvae from a hatchery experiencing larval mortality. While these strains caused minimal mortality, we categorize them here as “intermediate-avirulent” strains.

### 5.6 *V. mediterranei* comprises three phylogenetically distinct lineages

To understand the evolutionary basis for these phenotypic differences, we examined phylogenetic relationships among strains. We constructed a maximum likelihood phylogeny using 1,247 single-copy core genes from 33 *V. mediterranei* genomes, including our 11 experimental strains and 22 reference genomes from diverse marine sources. This analysis also included seven closely-related *V. barjaei* strains and two *V. thalassae* strains for phylogenetic context (Figure 5; Table 1). The phylogeny revealed three well-supported monophyletic clades that corresponded precisely with the experimentally determined phenotypes.

**Figure 5.**
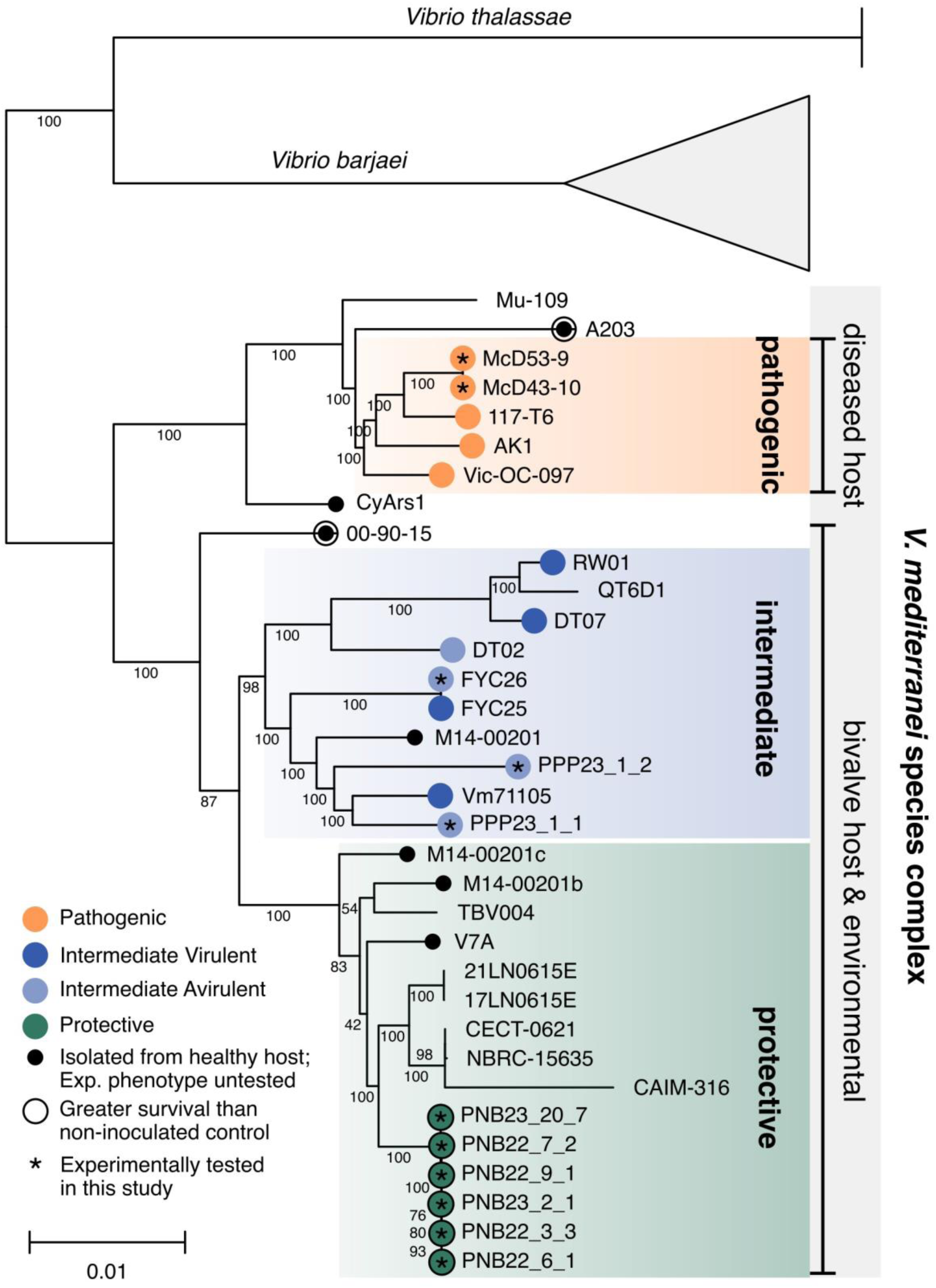
Phylogenetic analysis reveals three monophyletic clades within the *V. mediterranei* species complex. Maximum likelihood tree based on 1,247 single-copy core genes from 33 *V. mediterranei* genomes, plus representative *V. barjaei* and *V. thalassae* genomes. Bootstrap values shown at nodes. Colors indicate experimentally validated phenotypes from this study (stars) or isolation source and reported phenotypes from the literature (see Table 1). Clade designations (protective, pathogenic, intermediate) are based on strains with experimental phenotypic validation either in this study or previous literature. Pathogenic clade excludes strains isolated from healthy hosts (CyArs1, Mu-109, A203) despite phylogenetic proximity, as strain A203 enhanced *A. nauplii* survival when tested experimentally (Moreira et al. 2014), suggesting functional heterogeneity within this broader lineage. Strain 00-90-15 (black), isolated from healthy oysters and enhancing larval survival in experimental tests (Kehlet-Delgado, Häse, and Mueller 2020), falls outside defined clades. Three well-supported clades correspond to protective (teal, 100% bootstrap), intermediate (blue, 98% bootstrap), and pathogenic (orange, 100% bootstrap) lineages. Scale bar: 0.01 substitutions per site.

All six protective strains formed a tight monophyletic clade with 100% bootstrap support, designated the “protective clade.” This clade includes reference strain V7A from healthy *Argopecten purpuratus* (scallop) digestive tissue, which was previously reported to produce bacteriocin-like inhibitory substances and proposed as a potential probiotic (Serrano et al. 2018). Additional closely-related reference strains were isolated from seawater and sediment samples. While environmental reservoirs likely exist for all *V. mediterranei* lineages, the representation of isolates from the protective clade in seawater and sediment samples in particular suggests they persist outside host associations. The protective clade showed unexpectedly low internal genetic diversity, with all 15 strains from various sources sharing >99% average nucleotide identity (ANIm), consistent with recent shared ancestry or strong purifying selection maintaining beneficial traits.

The two pathogenic experimental strains clustered with reference strains isolated from diseased marine invertebrates, forming a “pathogenic clade” with 100% bootstrap support. This clade included *V. mediterranei* strains AK1 and Vic-OC-097, well-characterized coral pathogens, strain 117-T6 from diseased red algae (*Pyropia yezoensis*), and strains from moribund pen shells (*Pinna nobilis*). Within-clade ANIm values ranged from 98.5-99.9%, indicating somewhat greater genetic diversity than the protective clade while maintaining clear genomic cohesion.

An “intermediate clade” bridged the protective and pathogenic lineages, containing strains with variable virulence phenotypes from bivalve aquaculture systems and marine environments. This clade included our three intermediate-virulence experimental strains along with reference genomes from both healthy and diseased bivalves. The phylogenetic position of this clade, sister to the protective clade and divergent from the pathogenic lineage, indicates an intermediate genomic grouping consistent with their variable phenotypes.

Genomic divergence among clades approaches species-level thresholds while within-clade similarity remains high (Figure S1). ANIm analysis revealed consistently high within-clade values (Protective: 99-100%; Intermediate: 97.6-100%; Pathogenic: 98.2-100%), indicating recent shared ancestry or strong purifying selection. Between-clade comparisons revealed substantial genomic divergence, with Pathogenic-Intermediate strains showing 96.5-97.6% ANIm, Pathogenic-Protective showing 97.1-97.8% ANIm, and Intermediate-Protective showing 97.5-98.2% ANIm. These between-clade values approach the 95-96% ANI threshold typically used for species delineation (Richter and Rosselló-Móra 2009), highlighting substantial divergence within the species complex that is supported by their differing host-interaction phenotypes.

### 5.7 Protective and pathogenic lineages are distinguished by genomic specialization

Pangenome analysis revealed striking asymmetry in specialized gene content between clades. Protective strains harbored 230 unique orthogroups (genes present in 100% of protective strains and absent in all others), compared to 80 unique orthogroups in pathogenic strains and 27 in intermediate-virulent strains (Figure S2A; Table S1). Intermediate-avirulent strains notably lacked any clade-specific genes.

COG functional analysis of protective-specific genes identified enrichment in several categories: regulatory systems (21 transcriptional regulators including two-component systems EnvZ/OmpR), stress response mechanisms (27 DNA repair genes, nitric oxide dioxygenase), metabolic functions (glyoxylate carboligase, menaquinone biosynthesis pathway genes), and cell surface modification (19 cell wall biogenesis genes including multiple glycosyltransferases) (Figure S2B). Additional protective-specific elements included eight phage integrases and resolvases, Type IV secretion system components, and multiple toxin-antitoxin systems.

Among the protective-specific orthogroups, we identified an 88 amino acid predicted bacteriocin located within a putative biosynthetic operon containing genes encoding a peptide biosynthesis protein (YydG-like), acetyltransferase, and secretion machinery. This protein shares 45.6% identity with UniProt A0A240EP04, an annotated class II bacteriocin with double-glycine leader peptide in *Vibrio thalassae*.

Additional protective-specific orthogroups included genes involved in polysaccharide degradation (ß-1,3 glucan and glycogen metabolism), peptide transport systems, cell surface modification, biofilm formation, and pilus assembly.

The 80 pathogenic-specific orthogroups showed relatively limited functional diversity, distributed among secondary metabolism, transcriptional regulation, and amino acid/organic acid metabolism. The pathogenic clade uniquely encoded a complete Type I secretion system (T1SS) consisting of an ATP-binding protein (PrsD), membrane fusion protein (PrsE), and a large repetitive protein containing 44 hemolysin-type calcium-binding domains (1,480 amino acids). Additional pathogenic-specific functions included multidrug efflux pumps and specialized regulatory proteins.

Within the intermediate clade, virulence phenotypes correlated with the presence of specific genetic elements. All *V. mediterranei* strains regardless of clade were found to encode a type VI secretion system on chromosome I (T6SS1). However, intermediate-virulent strains uniquely possessed an additional T6SS on chromosome II (T6SS2), which was absent from both protective and pathogenic strains. Comparative analysis revealed that T6SS1 and T6SS2 are indeed structurally distinct systems, differing in both gene organization and amino acid identity of homologous core proteins (Figure S3).

Screening for the recently described *V. mediterranei* TCP pathogenicity island (TCP-PAI; (Zhang et al. 2025)) revealed this genomic region was absent from all protective strains. In contrast, TCP-PAI was variably present across intermediate and pathogenic clades, with considerable variation in gene content and completeness among strains where present. In protective strain Vm02, the genomic regions flanking the TCP-PAI insertion site are directly adjacent, confirming absence of the intervening pathogenicity island (Figure S4).

## 6 Discussion

Our findings establish that *V. mediterranei* comprises phylogenetically distinct lineages with fundamentally different host-interaction strategies, and we organize this discussion around four themes: the phylogenetic basis of protective versus pathogenic phenotypes, the genomic architecture underlying these strategies, implications for aquaculture under climate change, and broader considerations for *Vibrio* taxonomy and disease management.

### 6.1 Phylogenetically constrained protective versus pathogenic phenotypes in *V. mediterranei*

This study addresses the longstanding paradox of *V. mediterranei’s* contradictory roles across marine systems by demonstrating that beneficial and pathogenic phenotypes are phylogenetically constrained within distinct evolutionary lineages. The protective clade, comprising strains exclusively from healthy bivalves and environmental sources, forms a cohesive monophyletic group (>99% ANIm) that consistently rescued oyster larvae from near-complete pathogen-induced mortality. In contrast, pathogenic strains formed a separate lineage sharing only 97.1-97.8% ANIm with protective strains, approaching commonly cited species delineation thresholds (Richter and Rosselló-Móra 2009).

This phylogenetic structure reconciles contradictory literature: reports of *V. mediterranei* causing disease in corals (Kushmaro et al. 2001; Rubio-Portillo et al. 2018), pen shells (Andree et al. 2021), and razor clams (Fan et al. 2023) derive solely from strains within the pathogenic or intermediate clades. Conversely, strains described as commensal or beneficial – including the bacteriocin-producing V7A isolate from a scallop (Serrano et al. 2018) – cluster within the protective clade. The intermediate clade’s variable virulence, linking to presence or absence of T6SS2, illustrates how discrete genetic changes are associated with pathogenicity to a bivalve host. This mirrors observations in *V. fischeri* where different T6SS clusters contribute to niche partitioning and eukaryotic antagonism (Speare et al. 2018; Gaddy et al. 2025).

### 6.2 Genomic architecture of protection: Specialized defenses

The marked asymmetry in clade-specific gene content reveals fundamentally different evolutionary strategies, with protective lineages maintaining substantially greater genomic complexity than pathogenic strains (230 versus 80 unique orthogroups). This pattern suggests that stable beneficial associations require more specialized molecular machinery than opportunistic pathogenicity, paralleling other mutualistic systems where symbionts maintain expanded metabolic and regulatory repertoires to persist within host environments (Schwartzman and Ruby 2016; Soto and Nishiguchi 2021).

The protective clade’s genomic specialization includes unique oligopeptide transporters, carbohydrate utilization pathways (glyoxylate carboligase and glycogen metabolism), stress-response systems (nitric oxide dioxygenase, multiple DNA-repair genes), and notably, a predicted bacteriocin biosynthetic operon present in all protective strains. This combination suggests multiple candidate mechanisms that could contribute to protection, including: (1) competitive exclusion through efficient nutrient acquisition in the larval digestive environment; (2) direct antimicrobial activity against invading pathogens; (3) niche preemption through stable tissue colonization; and (4) enhanced stress tolerance enabling persistence under host immune pressure.

While our larval challenge assays demonstrate robust pathogen protection, the specific molecular mechanisms remain to be directly tested. The genomic architecture is consistent with protection operating through multiple complementary pathways rather than a single mechanism, aligning with emerging models of microbiome-mediated protection in which beneficial taxa simultaneously compete for resources, physically block pathogen colonization sites, and produce targeted antimicrobials (Buffie and Pamer 2013; Spragge et al. 2023). Functional validation of these candidate mechanisms, including bacteriocin activity, resource competition dynamics, and colonization resistance, represents an important avenue for future mechanistic studies.

In contrast, pathogenic lineages encode broad-spectrum offensive systems characteristically associated with anti-eukaryotic activity and tissue invasion. In the pathogenic lineage this includes a complete Type I secretion system (T1SS) and associated hemolysin-type calcium-binding domains. In the intermediate-virulent lineage this includes a second Type VI secretion system (T6SS2) absent from all other lineages. The protective clade’s complete absence of T1SS and T6SS2, combined with unique metabolic specialization and stress-tolerance pathways, is consistent with a model in which protective strains favor long-term niche occupation over host damage (Figure 6).

**Figure 6.**
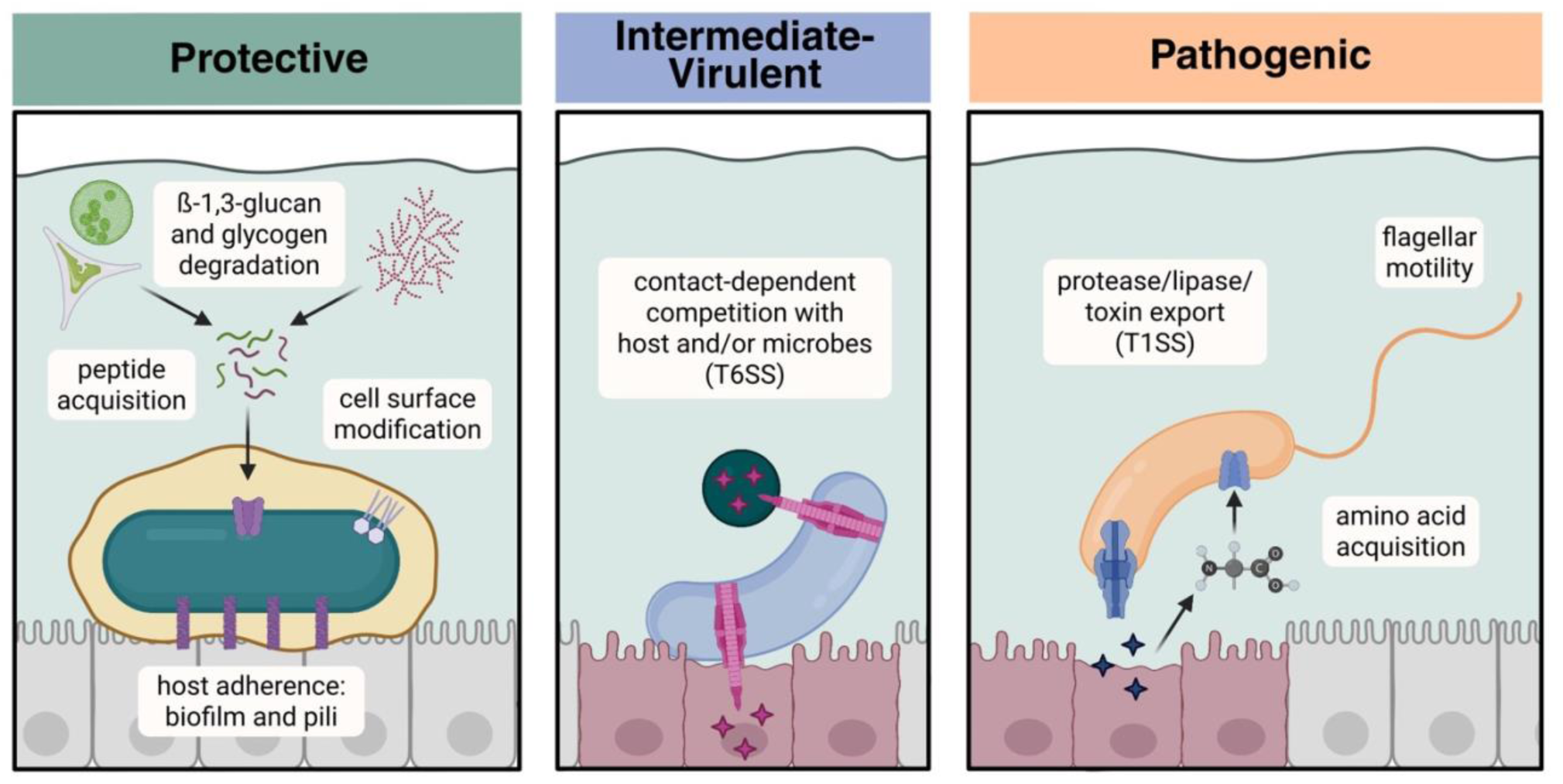
Proposed ecological strategies of *V. mediterranei* lineages. Schematic model illustrating distinct host-interaction mechanisms. Protective strains (left) colonize digestive tissues through metabolic specialization and stress tolerance. Intermediate strains (center) show variable virulence depending on T6SS2 presence. Pathogenic strains (right) deploy offensive secretion systems (T1SS) for tissue invasion.

### 6.3 Protection under thermal stress and climate change implications

The maintenance of protective efficacy at 32°C, despite increased virulence of both *V. harveyi* and *V. coralliilyticus* at elevated temperatures, has critical implications for aquaculture under warming ocean conditions. Marine heatwaves are increasing in frequency and duration (Oliver et al. 2018), and many *Vibrio* pathogens exhibit temperature-dependent virulence (Green et al. 2019; Vezzulli et al. 2016). Our finding that protection persists through 96+ hours across elevated temperatures suggests that protective *V. mediterranei* may serve as an effective biological control agent even under thermal stress conditions in shellfish hatcheries.

Fluorescence-based colonization assays demonstrate that at least one of the protective strains, Vm02, stably associates with larval digestive tissues, with this association established rapidly even when larvae are co-colonized with protective symbiont and pathogen simultaneously. This durability addresses long-standing challenges in probiotic development, where strain instability and inconsistent colonization frequently hamper efficacy (Kesarcodi-Watson et al. 2008).

### 6.4 Taxonomic implications and evolutionary dynamics

The discovery that *V. mediterranei* comprises deeply divergent protective and pathogenic lineages (97.1-97.8% ANIm) with mutually exclusive ecological strategies raises important considerations for *Vibrio* taxonomy. When *V. shiloi* was first described as a coral pathogen, it was explicitly distinguished from *V. mediterranei* on the basis of pathogenicity and cellular characteristics (Kushmaro et al. 2001). Our phylogeny recapitulates these distinctions: pathogenic isolates cluster with the original *V. shiloi* strains AK1 and Vic-OC-097, whereas protective and environmentally associated isolates group with the classical *V. mediterranei* lineage. The subsequent synonymization of these taxa (Thompson et al. 2001) may therefore have consolidated biologically heterogeneous lineages under a single name.

Regardless of formal nomenclature, our results underscore that species-level identification alone is insufficient for predicting ecological behavior within *V. mediterranei*. The clade-specific genomic markers identified here (e.g., bacteriocin operons in protective strains or T1SS/T6SS2 in pathogenic strains) provide practical tools for distinguishing beneficial from harmful populations in aquaculture settings and for interpreting historical reports of “*V. mediterranei*” in the literature.

The near-clonal structure of the protective clade across years and sources (>99% ANIm) contrasts with greater genomic diversity in the pathogenic lineage (98.5-99.9% ANIm), potentially reflecting broader ecological opportunity or adaptation to diverse hosts. The intermediate clade’s variable virulence, defined by presence or absence of T6SS2, highlights how mobile genomic elements may enable rapid shifts between commensal and pathogenic lifestyles (Bruto et al. 2018; Lemire et al. 2015).

These findings suggest new monitoring approaches that extend beyond traditional pathogen surveillance. Our recent field study showed systematic depletion of protective *V. mediterranei* preceding mortality events (Smith et al. 2025), indicating that protective taxa may function as early-warning biomarkers of community disruption. Clade-specific dPCR assays targeting the protective lineage’s unique genes could provide high-resolution tracking of beneficial *Vibrio* populations, while assays detecting T1SS or T6SS2 may allow hatcheries to identify emerging threats before clinical disease becomes apparent.

### 6.5 Limitations and future directions

Several limitations should be considered. Our larval model provides proof-of-concept but may not fully represent interactions in adult oysters or in complex field environments. Additionally, our experiments assess protection only through the pre-metamorphosis larval stage (up to 11 days post-fertilization, 96 hours post-bacterial exposure). Whether protective *V. mediterranei* can persist through metamorphosis and continue providing benefits in juvenile or adult oysters remains unknown but is critical for practical probiotic deployment.

Community-level processes, such as polymicrobial interactions and host genotype-by-microbe effects, remain to be explored. While we cannot exclude geographic sampling bias, strains from each clade derive from diverse geographic locations (protective strains from North Carolina, Australia, Chile, and the Mediterranean; pathogenic strains from Florida and the Mediterranean), suggesting that phenotype rather than geography drives phylogenetic clustering. Notably, colonization capacity was directly demonstrated for strain Vm02. The high genomic similarity within the protective clade (>99% ANIm) raises the hypothesis that colonization may be a shared trait of this lineage, but this requires experimental validation across additional strains. While genomic analyses suggest candidate protection mechanisms involving metabolic competition, antimicrobial activity, and enhanced stress tolerance, direct functional validation of these mechanisms is still needed. Finally, the evolutionary stability of beneficial phenotypes requires assessment. The protective clade’s monophyletic structure, extensive genomic investment in specialized functions, and lack of T1SS and T6SS2 suggest stable mutualistic traits. However, the intermediate clade demonstrates that horizontally acquired elements can rapidly alter ecological strategy. Long-term monitoring of protective strain genomic stability, particularly under different selection pressures, will be important for ensuring reliable probiotic performance.

### 6.6 Conclusions

This study redefines *V. mediterranei* as a species complex comprising lineages with fundamentally distinct ecological strategies. Beneficial and pathogenic phenotypes correspond to phylogenetically constrained clades rather than plastic responses to environmental conditions. Identification of a monophyletic protective clade that confers robust pathogen protection under thermal stress conditions challenges pathogen-centric views of disease ecology and underscores the importance of preserving beneficial microbiome members in aquaculture systems. As marine ecosystems face increasing thermal and disease stress, recognizing and maintaining protective symbionts like *V. mediterranei* will be as critical as controlling pathogens. By combining field observations, experimental validation, and comparative genomics, we present a framework for uncovering cryptic beneficial relationships and highlight new opportunities for sustainable disease management in marine hatcheries.

## 9 Author Contributions

Steph Smith: Conceptualization, Methodology, Investigation, Formal Analysis, Writing – Original Draft, Visualization, Project Administration

Ami Wilbur: Resources, Methodology, Writing – Review & Editing Jaypee Samson: Investigation, Resources

Yesmarie De La Flor: Investigation, Resources

Marta Gomez-Chiarri: Resources, Methodology, Writing – Review & Editing Blake Ushijima: Resources, Methodology, Writing – Review & Editing

Rachel Noble: Conceptualization, Funding Acquisition, Supervision, Writing – Review & Editing All authors read and approved the final manuscript.

## 11 Acknowledgments

We thank the UNCW Shellfish Research Hatchery for providing larval oysters and hatchery facilities essential to this study. We are grateful to Tal Ben-Horin for providing access to fluorescence microscopy equipment and expertise on oyster physiology, and to Thomas Clerkin for extensive feedback on experimental design. The *V. mediterranei* strains McD53-9 and McD43-10 were originally isolated at the Smithsonian Marine Station at Fort Pierce and were graciously provided by Dr. Valerie Paul.

## 12 Data Availability Statement

Raw sequencing reads for the newly sequenced genomes (McD43-10, McD53-9, FYC25, FYC26, PPP23_1_1, PPP23_1_2) have been deposited in the NCBI Sequence Read Archive under BioProject accession PRJNA1379678. Assembled genomes have been deposited in NCBI GenBank with accession numbers provided in Table 1.

## Supplemental Information

**Figure S1.**
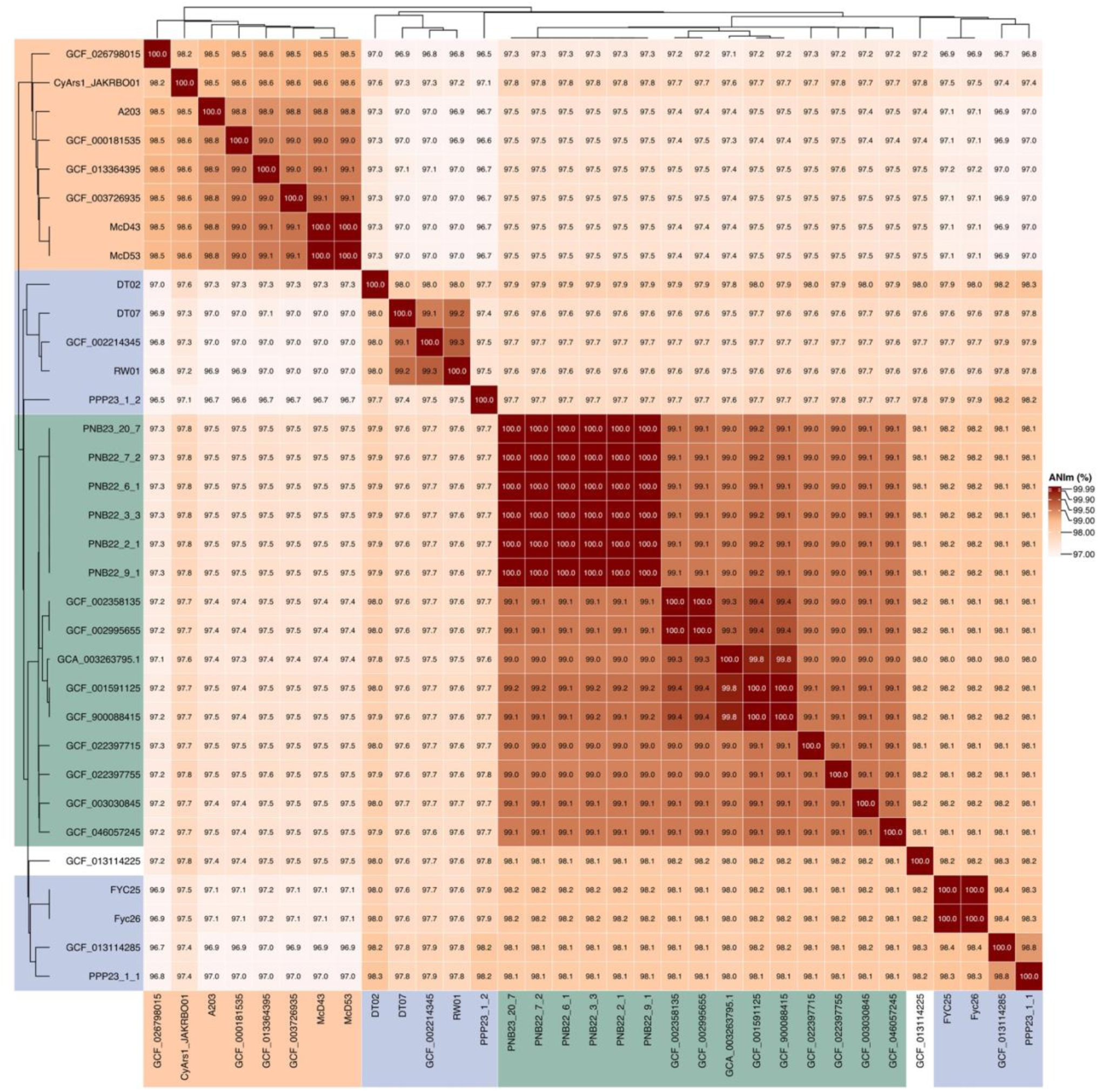
Genomic divergence within *V. mediterranei* revealed by average nucleotide identity. Heatmap of pairwise ANIm values for 33 genomes. Within-clade identity: Protective 99-100%, Intermediate 97.6-100%, Pathogenic 98.2-100%. Between-clade identity: 96.5-97.8%, approaching species boundary threshold (95-96%).

**Figure S2.**
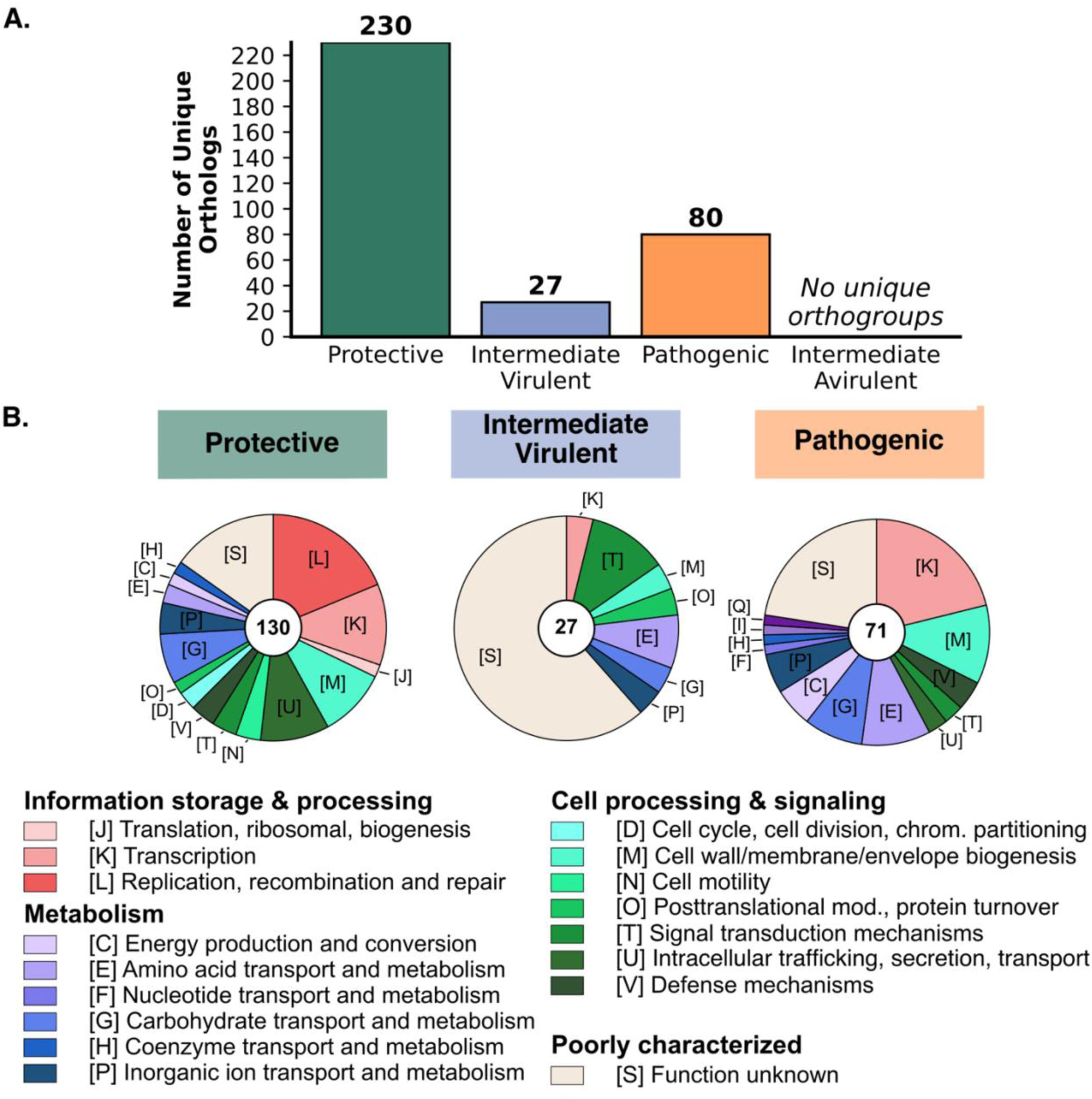
Functional specialization of phenotype-specific genes. **(A)** Asymmetric distribution of unique orthogroups: protective strains harbor 230 unique genes versus 17 in pathogenic strains. **(B)** COG functional distribution of phenotype-specific genes.

**Figure S3.**
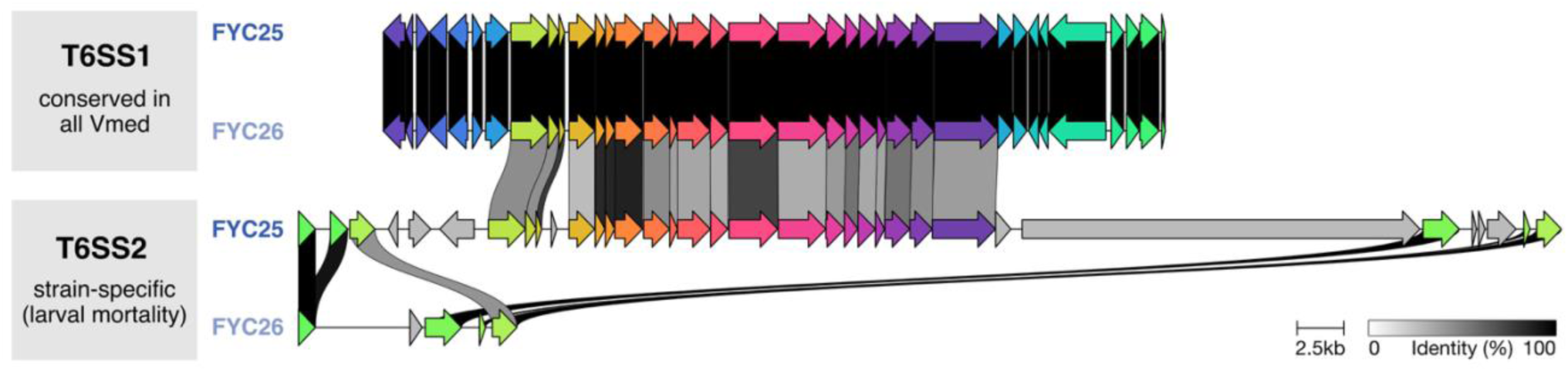
Type VI secretion system distribution in intermediate *V. mediterranei* strains. Comparative genomic alignment showing T6SS1 (conserved in all strains regardless of phenotype) and strain-specific T6SS2 presence in virulent intermediate strain FYC25 but absence in avirulent FYC26. Identity scale and gene clusters shown.

**Figure S4.**
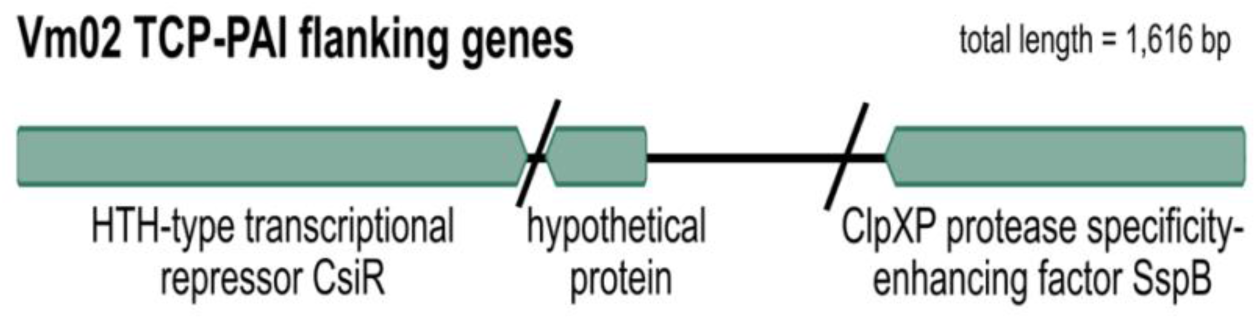
TCP pathogenicity island absent from protective *V. mediterranei* strains. Genomic context of TCP-PAI flanking region in protective strain Vm02 showing direct junction of flanking genes (1,616 bp total) without intervening pathogenicity island.

