## Supplemental Information (Figs S1-S4) for "Phylogenetically distinct *Vibrio mediterranei* lineages confer robust protection under thermal stress against oyster pathogens"

### SUPPLEMENTAL MATERIALS

#### SUPPLEMENTAL FIGURES

##### Figure S1

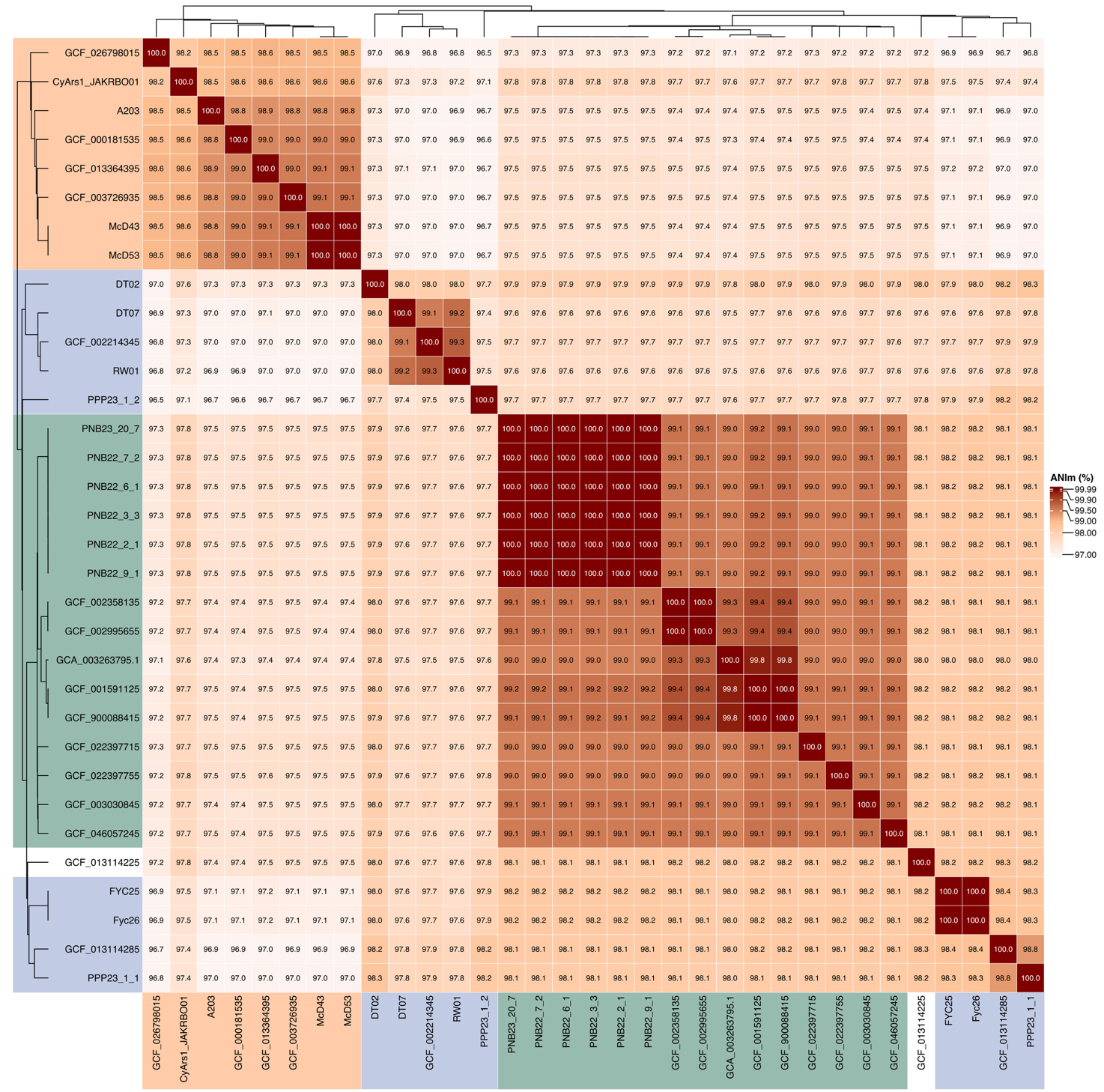

##### Figure S1. Genomic divergence within *V. mediterranei* revealed by average nucleotide identity. Heatmap of pairwise ANIm values for 33 genomes. Within-clade identity: Protective 99-100%, Intermediate 97.6-100%, Pathogenic 98.2-100%. Between-clade identity: 96.5-97.8%, approaching species boundary threshold (95-96%).

##### Figure S2

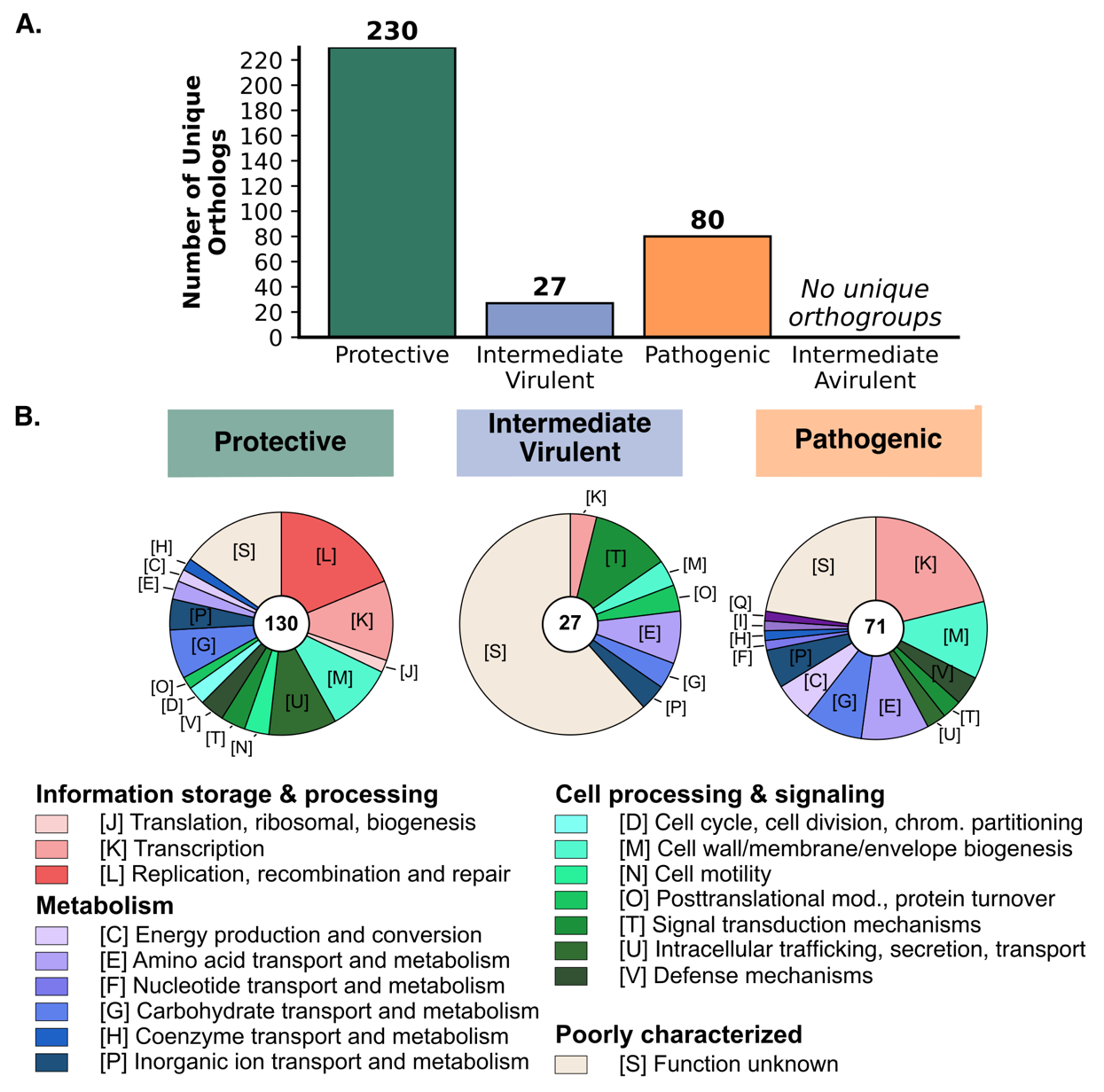

##### Figure S2. Functional specialization of phenotype-specific genes. (A) Asymmetric distribution of unique orthogroups: protective strains harbor 230 unique genes versus 17 in pathogenic strains. (B) COG functional distribution of phenotype-specific genes.

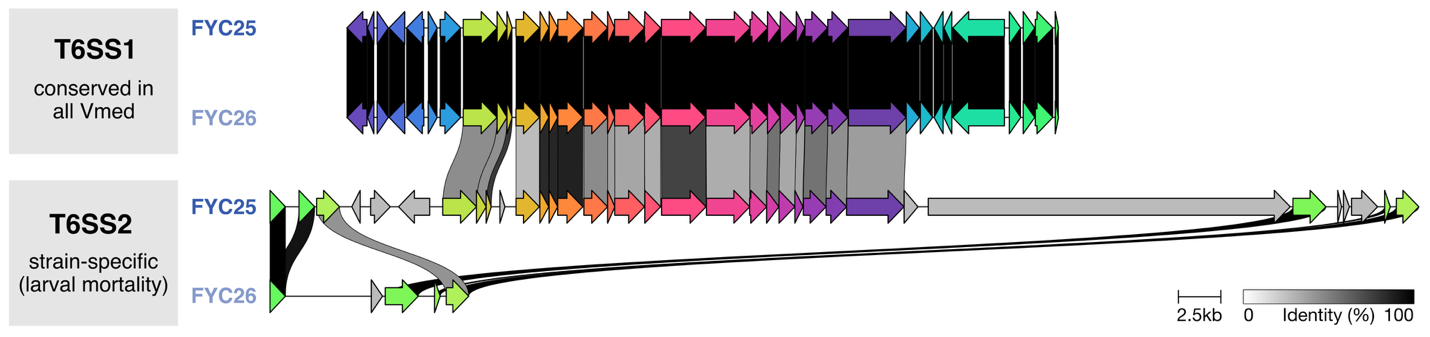

##### Figure S3. Type VI secretion system distribution in intermediate *V. mediterranei* strains. Comparative genomic alignment showing T6SS1 (conserved in all strains regardless of phenotype) and strain-specific T6SS2 presence in virulent intermediate strain FYC25 but absence in avirulent FYC26. Identity scale and gene clusters shown.

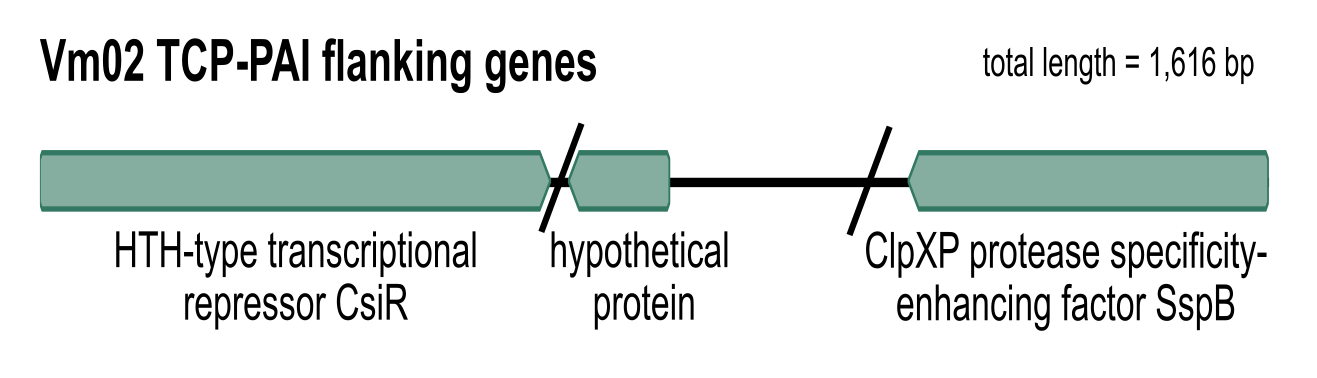

##### Figure S4. TCP pathogenicity island absent from protective *V. mediterranei* strains. Genomic context of TCP-PAI flanking region in protective strain Vm02 showing direct junction of flanking genes (1,616 bp total) without intervening pathogenicity island.
